# Construction, Phenotypic Characterization, and Immunomodulatory Function Study of BMSC-Macrophage Hybrid in vitro

**DOI:** 10.1101/2025.07.25.666751

**Authors:** Jian Shi, Xiaotong Wu, Anqi Yang, Jiawei Pan, Yangyang Sun, Jundong Zhu, Yuan Zhang, Linglong Jiang, Min Fan

## Abstract

**Objective:** The aim of our study was to integrate bone marrow-derived stem cells (BMSCs) with macrophages and to investigate the performance of this hybrid in IgG clearance, immunosuppression, and the mitigation of inflammatory injury.

**Methods:** BMSCs and RAW264.7 cells were fused through transient transfection using the COVID-19 spike glycoprotein and human angiotensin-converting enzyme 2 (hACE2), respectively. The resulting hybrids were purified using magnetic sorting, and their characteristics were evaluated by examining morphology, glucose uptake rates, phagocytic capacity, cytoskeletal morphology, and the levels of Fcγ receptors, as well as anti-inflammatory, antioxidant, adhesion, and complement inhibitory factors. The IgG clearance capability of the hybrids was assessed by measuring internalization efficiency and infiltration into renal organoids. Additionally, the reparative effects of the hybrids were evaluated using doxorubicin-treated MPC5 damage models. Live cell imaging techniques were employed to investigate the interactions between the hybrids and immature dendritic cells (iDCs). Hybrid-derived nanovesicles were isolated, and their abilities to target IgG and clear IL-6 were evaluated using immunofluorescence and ELISA. Furthermore, chloroplasts and C-dots were co-incubated with the hybrids, and their benefits were assessed by measuring cell viability, oxygen content, ROS levels, mitochondrial function, and the levels of anti-inflammatory, antioxidant, and pro-repair factors. A scratch assay was conducted to determine the impact of chloroplast transplantation on the migration ability of 3T3 cells.

**Results:** High-purity hybrids were successfully produced that cleared IgG while maintaining their anti-inflammatory and antioxidant effects. They promoted the recovery of podocyte injury through vesicular and mitochondrial delivery, as well as efferocytosis. Furthermore, the hybrids inhibited iDCs primarily through migrasomes, intercellular nanomicrotubule connections, and phagocytosis. The derived nanovesicles were capable of residing in IgG-enriched regions and adsorbing IL-6. Interestingly, the transplanted chloroplasts enabled the hybrids to utilize light energy to enhance their antioxidant capacity and promote the migration of 3T3 cells, which contributes to wound repair. Subsequently, the loading of C-dots was beneficial for enhancing resistance to oxidative damage.

**Conclusions:** Our results suggest that hybrids-mediated therapy is a new and creative therapeutic approach for managing immune-mediated nephropathies.Hybrids demonstrated effective immunomodulatory and promoted injury recovery. The derived nanovesicles have the potential to alleviate the inflammatory burden at sites of antibody immune complex deposition. Additionally, the incorporation of chloroplasts and C-dots enhanced the hybrids’ adaptability to ischemic-hypoxic microenvironments.

## Introduction

Immunonephropathy is a chronic glomerular disease characterized by shared immunopathologic features across multiple etiologies, including purpura nephritis, lupus nephritis, and IgA nephropathy, which arise from immune system dysfunction^1–3^. This dysregulation activates immune cells, leading to the production of antibody-mediated immune complexes that deposit in renal tissues. These deposits induce intrinsic renal cell damage, trigger inflammatory cascades, and disrupt normal cellular functions, culminating in nephropathic manifestations such as proteinuria, hematuria, edema, and progressive renal impairment^4^. Notably, approximately one-third of patients eventually develop uremia. Current therapeutic strategies, including plasma exchange, immunosuppressants, corticosteroids, and antiplatelet agents, aim to alleviate symptoms and delay disease progression^5^; however, their efficacy remains suboptimal, underscoring the need for novel interventions.

In the renal microenvironment of immune nephropathy, extensive macrophage infiltration occurs, with M1 macrophages secreting pro-inflammatory factors that exacerbate the disease. In the later stages, although M2 anti-inflammatory macrophages are also activated, they secrete pro-fibrotic factors that promote renal fibrosis^6,7^. Notably, macrophages exhibit a degree of phagocytosis of immune complexes. Additionally, they mediate antibody-dependent cell-mediated cytotoxicity (ADCC), up-regulate pro-inflammatory factors, and contribute to kidney damage. While many researchers have recently focused on utilizing M2-type macrophages for the treatment of immune nephropathy, these macrophages are prone to polarization driven by the inflammatory microenvironment, resulting in a limited therapeutic effect^8^.

BMSCs have been reported to prevent kidney injury and promote renal recovery by modulating the immune system through paracrine effects^9^. Nevertheless, the therapeutic effects of BMSCs in vivo are time-limited and suboptimal for treating severe or chronic immune diseases. Furthermore, the phagocytic capacity of BMSCs is limited, making the removal of immune complexes a significant challenge.

Interactions between BMSCs and macrophages have been observed, with evidence demonstrating that BMSCs can restore the diaphragm injury gap between peduncles by reducing macrophage infiltration and polarizing them towards the M2 phenotype^10^. Additionally, BMSCs has been shown to promote the release of anti-inflammatory factors, reduce the release of pro-inflammatory factors, and enhance phagocytosis of macrophage. Moreover, the anti-inflammatory, pro-repair, pro-angiogenic, and immunosuppressive functions of BMSCs are further enhanced by macrophages^11^. However, clinical findings indicate that the proportion of BMSC-macrophage interactions in vivo is minimal and predominantly occurs via paracrine secretion^12^. Therapeutic methodologies aimed at enhancing these interactions demonstrate limited efficacy. Consequently, we propose the hypothesis that fusing BMSCs and macrophages may leverage their respective strengths while mitigating their weaknesses through their interactions, thereby amplifying the anti-inflammatory and immunosuppressive effects. This fusion could potentially serve as a novel treatment for immune nephropathies.

Cell fusion is defined as the process of merging two or more cells into a single cell, either under natural conditions or through artificially induced methods^13^. This process encompasses the connection and fusion of plasma membranes, the merging of cytoplasm, and the mutual formation of nuclei, organelles, and enzymes into a composite system. The resultant hybrid, formed through cell fusion, exhibits a variety of cellular characteristics, with certain cellular functions being amplified and enhanced. The occurrence of cell fusion has been demonstrated within the tumor microenvironment and the inflammatory immune microenvironment^14^. Existing induction techniques, including PEG, electrofusion, and co-incubation, have shown low levels of fusion efficiency and specificity, thereby limiting their clinical applicability^15–17^.

In this study, novel methods were explored for the induction and purification of macrophages and BMSCs cells to form hybrids in vitro, aiming to investigate their immunomodulatory role in promoting new therapies for clinical immune nephropathies. Furthermore, the employment of xenografting of cellular organelles (chloroplasts) and carbon quantum dot loading was attempted, with the objective of enhancing the therapeutic function of hybrids.

## Materials and Methods

### Cell Culture and 3D Kidney Organoid Construction

BMSC were sourced from Ruyaobio, while the cell lines RAW264.7, MPC5, DC, and PC3 originated from MeisenCTCC. The BMSC were grown in DMEM/F12 medium(Gibco,21127022) enhanced with 10% fetal bovine serum (FBS, VivaCell, C04001) and 1% streptomycin-penicillin (SP, Gibco, 15240062). RAW264.7 cells thrived in DMEM medium(Gibco, 11995065) containing 10% FBS and 1% SP. Meanwhile, the MPC5 cells were nurtured in DMEM medium supplemented with 10% FBS and 1% SP. In the case of iDC cells with inherent green fluorescence, they were cultured in IMDM medium(Gibco, 31980030) that included 10% FBS and 1% SP. MPC5 cells were specifically cultivated in 12-well plates. Following a 24h treatment with 0.5 μM Doxorubicin (DOX,MCE, HY-15142), various analyses focused on apoptosis, JC-1, ROS, and CD80 expression were performed utilizing a flow cytometer, along with qRT-PCR for measuring IFNγ, IL-6, TNF-α, and IL-1β levels. To further develop the inflammatory injury model of kidney organoids, MPC5 cells underwent sequential labeling with antiCD80(Abcom,ab222720) and mouse anti-Rabbit IgG-CoraLite488(secondary antibody, Proteintech, SA00014) followed by exposure to DOX in a 2D well plate. The collected cells were then transformed into 3D microsphere organoids utilizing NAC-Organ technology, which is based on nano-nucleic acid material (PUHENG Bio).

### Cell Fusion, Purity and Characterization

BMSC and RAW264.7 cells were cultured to 70% confluence prior to plasmid transfection. Specifically, BMSC utilized plasmid COVID-19-S(purchased from Honor Biotech), while RAW264.7 employed plasmid hACE2(purchased from Honor Biotech). Following 24h of transfection, BMSC and RAW264.7 cells were mixed in equal proportions and incubated for an additional 24h to promote spontaneous fusion, after which unattached cells were removed. To symbolize the hybrids, PKH67-labeled BMSC and PKH26-labeled RAW264.7 cells were employed to detect membrane fusion. Additionally, Mito-Green tracker(Dojindo, MT10)-stained BMSC and Mito-Red tracker(Dojindo, MT11)-stained RAW264.7 cells were utilized to observe cytoplasmic fusion, following the previous instructions. DiD-labeled BMSCs and PKH26-labeled RAW264.7 cells were mixed, and the Mito-Green tracker was employed to distinguish the mitochondrial populations between the hybrid cells and BMSCs under flow cytometry, without the presence of visible RAW264.7 cells. To obtain pure hybrids, the mixed cells were labeled with 15 µg/ml of superparamagnetic nanoparticles(Nanoeast, Mag3200-050). The hybrids were subsequently extracted using a directional static magnetic field within 30 min, and the purity of the hybrids was determined by assessing the percentage of cells that were positive for both DiD and DiO, as described above. Moreover, to evaluate the morphology of cells, scanning electron microscopy (SEM) was employed. In the SEM protocol, cells were fixed in 4% paraformaldehyde (PFA) for 45 minutes at room temperature. This was followed by washing with distilled water and dehydrating through a stepwise ethanol series. Subsequently, the samples were coated with gold using a ScanCoat 6 (UK) and analyzed with a SE1 detector configured at a filament voltage of 1 kV. Hybrid cells along with BMSCs were rinsed using PBS and then incubated in serum-free, sugar-free DMEM that was supplemented with 2-NBDG reagent (Thermo, N13195) at a final concentration of 200 µg/mL for a duration of 1h. After the incubation, the cells were collected from the plate and washed with PBS. Glucose uptake evaluations were conducted immediately via flow cytometry analysis following the manufacturer’s guidelines.

### Prussian blue Dyeing

The adherent cells were removed from the culture medium and subsequently washed twice with 1× PBS. They were then fixed in 4% PFA for 15 minutes. Following fixation, the samples were treated with freshly prepared Perls staining solution(Solarbio, G1426) incubated at 37°C for 30min, and washed in distilled water for 2 min. Afterward, the samples were treated with Perls re-staining solution and re-stained for 1 minute. Finally, the samples were rinsed with water and examined under a microscope.

### JC-1、ROS and Apoptosis Assay

The assessment of mitochondrial membrane potential was conducted using JC-1 (Solaibao, M8650). In a 12-well plate, 0.5 mL of medium was combined with 0.5 mL of the JC-1 working solution prepared and added to the cells. After incubating for 30 minutes at 37°C in darkness, the cell suspensions underwent two washes with JC-1 staining buffer. The resulting cell pellets were then resuspended in medium and analyzed via flow cytometry. Cultured cells were stained with the CellROX probe (Maokangbio, C10422) or intracellular ROS detection kit(Beyotime, S0033S) for 30 minutes at 37℃. Following washing, they were promptly collected and measured using flow cytometry. An apoptosis detection kit (Beyotime, C1062S) was employed to evaluate apoptosis. After treatment, cells underwent washing with PBS and were then resuspended in 250µl of binding buffer. Next, 5µl PI and 2.5µl AnnexinV-FITC were introduced and mixed well. The mixture was incubated in the dark at room temperature for 5 min. Ultimately, the rate of apoptosis was assessed using a flow cytometer.

### Colony Formation Assay

The proliferative capacity of RAW264.7, BMSC and hybrid cells was assessed using a plate cloning assay. Cell suspensions containing 1000 cells per well were seeded into 6-well plates, followed by continuous cultivation for 7 days. The visible cell colonies were fixed with 4% PFA and stained with a crystal violet solution (Beyotime, C0121) for 10 minutes. Typical images of the colonies were captured using a camera and a microscopic imaging system.

### Human Erythrocyte Fragment Phagocytosis Assay

In the experiment assessing the in vitro phagocytic capacity, a volume of 2 ml of human anticoagulated blood was utilized to enrich erythrocytes via lymphocyte separation medium. Following the staining of erythrocytes with DiO, lysis was performed using sterile water. The resulting pellets were harvested and co-cultured with RAW264.7, BMSCs, and hybrids for a duration of 4h. Flow cytometry was subsequently employed to measure uptake efficiency, relying on fluorescence intensity for assessment.

IgG Phagocytosis Test

20 μg of IgG-FITC antibody(Tonbo, 35-4724-U100) was incubated with hybrids for a duration of 30 min. After this period, the supernatant was removed, and the cells underwent three washes with PBS to facilitate the observation of antibody localization within the cells. Next, the experiment consisted of three distinct groups: ⍰ hybrid, ⍰ hybrid + IgG, and ⍰ (IFNγ/IL-1β) hybrid + IgG. For group ⍰, a pretreatment with IFNγ/IL-1β was applied for 24h, using a concentration of 20 ng/ml for each cytokine. After this treatment phase, flow cytometry analysis was conducted on all three groups.

To evaluate the expression of FcγR, as well as anti-inflammatory and antioxidant factors, adhesion molecules, pro-inflammatory agents, and complement inhibitory-related proteins in hybrids following IgG phagocytosis for 24h, qRT-PCR was performed.

### ADCC Detect

MPC5 cells were stained with cell tracker deep red(Maokangbio, MX4110), followed by the addition of an equal quantity of hybrids for a 24h co-culture. After this period, both the supernatant and the adherent cells were gathered to evaluate the apoptosis rate of MPC5 cells using a PI staining kit(Beyotime, C1062S) and analyzed by flow cytometry.

### MPC5 and Hybrid Interaction Pathway Assay

MPC5 cells were treated with DOX and subsequently labeled with antibodies as outlined in previous studies. Following this, they were co-cultured with hybrids for 24 hours to examine morphological changes in the MPC5 cells using a bright field microscope. Afterward, the MPC5 cells were labeled with cell tracker deeep red, and their membranes were stained using the DiO probe. Next, primary and secondary antibodies against CD80 were introduced. The evaluation of the transmission through the MPC5 cell membrane was performed using flow cytometry post co-culture with the hybrid. In a corresponding experiment, the membranes of the hybrids were also labeled with the DiO probe, and flow cytometry was utilized to assess the transmission of the hybrid’s membrane. Furthermore, the hybrids’ mitochondria were stained with the Mito Green tracker, enabling the identification of mitochondrial transfer from hybrids via flow cytometry.

### ACs Preparation and Efferocytosis Test

MPC5 cells were gathered and dispersed in PBS to generate ACs through heating at 100°C for 15 minutes. The ratio of apoptotic cells in the pellets was measured using flow cytometry, adhering to previously established protocols. Following this, ACs were treated with PKH67 and co-cultured with hybrids at a 5:1 ratio for 4 hours before observation, as documented^18^. In addition, ACs were labeled with PKH26 and co-cultured with hybrids at the same 5:1 ratio for only 2 hours. Subsequently, the PKH67-stained ACs were introduced and incubated for another 2h prior to observation. To assess proliferation, hybrids were stained with CytoTell^TM^ Red 650(AAT, 22256) and co-cultured with non-labeled ACs for 4h, followed by three washes to eliminate any non-internalized ACs. The culture then continued for 24h before being analyzed by flow cytometry. Additionally, the phenotype of hybrids was assessed using qRT-PCR.

### Hybrid and iDC Interaction Measurement

The interaction between DiD-labeled hybrids and iDCs was studied using a long-term real-time dynamic live cell imaging analyzer (Essen BioScience, Inc.). The dynamic process was recorded over a period of 30 minutes to observe changes over 24 hours. Furthermore, hybrids pretreated with IFN-γ and IL-1β (20μg/ml for each) for 24h were cultured alongside iDCs, which were subsequently stained with CytoTell^TM^ Red 650 to assess the proliferation capacity of iDCs after 24 hours, following the reagent instructions. Additionally, after co-culture, the cells were harvested and resuspended in 100 μL of PBS, followed by the addition of 0.5 μL of PE-conjugated-PDL-1 antibody (eBioscience, 12-5982-81) to each sample, and incubated in the dark for 30 minutes. After washing, the PDL-1 levels on iDCs were analyzed using flow cytometry. The hybrid membranes were tagged with DiD and subsequently co-cultured with iDCs for a duration of 24h. Flow cytometry was then used to evaluate the percentage of iDCs that took up the hybrid cell membranes, along with any alterations in PDL-1 expression in the iDCs. To clarify the mechanism of membrane transfer between the two cell types, the hybrids were treated with nanotube inhibitors (L-778123, L77, MCE, HY-16273) and migrasome inhibitors (Dynasore, Dy, MCE HY-15304 and CK636, MCE HY-15892) for 8 hours prior to being co-cultured with iDCs for another 24 hours. Flow cytometry was utilized to assess both the uptake of the hybrid cell membranes by iDCs and the expression levels of PDL-1.

### M1 Type Macrophages Induction

Add LPS (5 ng/ml) and IFNγ (20 ng/ml), and incubate RAW264.7 and hybrid cells for 24h. Subsequently, the morphological changes of the cells were observed under a microscope, and the mRNA levels of CD206, Arg1, IL-10, iNOS, TNF-α, and IL-6 were analyzed to assess the polarized status of macrophages.

### Cell-Derived Vesicles Production

The hybrids were pelleted and resuspended in PBS at a concentration of 1-5×10^6 cells/ml. Approximately 1 ml of the suspension was placed in a nitrogen cavitation chamber to disrupt the cells. Following this, the mixture was centrifuged at 1000 g for 15 minutes to remove cell debris, and subsequently centrifuged at 3000g for 10 minutes to obtain hybrid membrane-formed nanovesicles (NVs). The sizes and morphological characteristics of the NVs were measured using Nanoparticle Tracking Analysis (NTA) and Cryo-TEM.

### Western Blot Analysis

Samples that contained hybrid-NVs were subjected to denaturation and then applied to a 10% PAGE gel. Following this, the proteins were transferred to polyvinylidene fluoride (PVDF) membranes. The membranes were blocked with a 5% BSA solution at room temperature for 1 hour and then incubated with primary antibodies: CD16, CD64, PDL-1, and IL-6R (SAB,56554; SantaCruz, sc-515431; SantaCruz, sc-518027; SAB, 41912) overnight at 4 °C. Lastly, the membranes were treated with a suitable HRP-conjugated secondary antibody (Abcom, ab205718, ab6728, ab47827) for 1 hour, and the blots were visualized using ECL reagent based on the intensity of the signals.

### Nanovesicles Label PC3 Cells

Hybrids were treated with the DiO dye, and afterward, the NVs were prepared according to the previously described protocol. PC3 cells were seeded onto a 12-well plate and fixed using an immunostaining fixative for 15 min. Following this, the cells underwent three washes with PBS. To enhance cell permeability, an immunostaining permeabilization solution was applied for 1 hour, succeeded by an additional three washes with PBS. The cells were then treated with 2% BSA for blocking 30 min, and were subsequently categorized into three distinct groups: ⍰ PC3 + mouse anti-human PSA antibody, ⍰ PC3 + NV, and ⍰ PC3 + mouse anti-human PSA antibody + NV. The PSA primary antibody was introduced to groups ⍰ and ⍰ for overnight incubation, which was followed by three PBS washes. For group ⍰, a secondary antibody was incorporated for 1 hour to evaluate the impact of PSA antibody labeling. After this incubation period, NVs were added to groups ⍰ and ⍰ for co-incubation lasting 2h, followed by three washes with PBS. The samples were examined using a fluorescence microscope.

### ELISA Assay for Mouse IL-6

IL-6 concentration was carried out using an ELISA kit(Youkelife, EM0004). The cells were categorized into four distinct groups: ⍰ PC3 + IL-6; ⍰ PC3 + IL-6 + PSA antibody; ⍰ PC3 + IL-6 + NV; ⍰ PC3 + IL-6 + PSA antibody + NV. Following a 2h incubation with mouse IL-6, the supernatant was collected. Subsequently, the samples underwent centrifugation at 1200 rpm for 10 minutes, after which the supernatant was harvested for IL-6 analysis according to the procedure.

### Chloroplast Extraction, Cell Loading and Relative cpDNA Content Assay

Chloroplasts were isolated from young spinach leaves according to the procedures outlined in the operation manual^19^. The extracted chloroplasts were resuspended in 1 ml of PBS, and subsequently, 5 µl of this suspension was added to the hybrid cultured in a 12-well plate. Subsequent 24-hour light treatment for the hybrids was conducted using household LED full-spectrum plant grow light(WEN-1) purchased from Guixiang Company, with a wavelength range of 400-800 nm and an intensity of 50 µmol photons m^−2^s^−1^. Chloroplast uptake was monitored using a live cell dynamic detector within 3 h, while retention time was assessed with a microscope after 24h of incubation. To verify the chloroplast internalization pathway, the hybrid was pretreated with Dynasore for 8h prior to the addition of chloroplasts. The number of chloroplasts present in the cells was then evaluated after an additional 24h. The DNA of chloroplasts in hybrids was extracted from the total cellular DNA using a DNA extraction kit(Vazyme, DC112). Subsequently, qRT-PCR was employed to detect the Ct values and melt curves of the Dloop (a mitochondrial marker) and PsbA (a chloroplast marker).

### Intracellular O_2_ Level Evaluation

Hybrids were cultured under 1% O_2_ in a starvation medium containing 20% complete medium and 80% EBSS for 24h to simulate the unfavorable environments encountered during stem cells treatment for skin wounds. Subsequently, the hybrids were stained with a 5μM [Ru(dpp)_3_]Cl_2_ probe (MaoKangbio, MX4826) for 15 minutes, and the fluorescence intensity was analyzed using flow cytometry.

### Chloroplast Transmission Assay

Hybrids containing DiO-labelled chloroplasts were co-cultured with 3T3 cells in a transwell chamber for 24h in 12-well plates. Subsequently, the 3T3 cells were collected to measure the DiO-positive cell rate using flow cytometry.

### Wound Healing Assay

3T3 were cultured in 12-well plates and subsequently scratched using a sterile 200 µL pipette tip. Following the wounding, the above cells were co-cultured hybrids loading chloroplast using transwell chamber under starvation medium as previous described with 1%O_2_ in the incubator. After 24 hours, the migration of 3T3 cells was examined by microscope.

### Synthesis and Identification of C-Dots

The C-dots were manufactured using a streamlined one-pot solvothermal method as reported previously^20^. In summary, 1.2 g of D-biotin and 1.2 g of GSH were combined with 40 ml of formamide. The resulting solution was then transferred to a Teflon autoclave and heated at 160 °C for 8 hours. After cooling, the products were dialyzed in deionized water (molecular weight cut-off (MWCO) 3500 Da) for 7 days. Subsequently, the products were filtered using 0.22 μm filters, and the process concluded with concentration and lyophilization. The visible absorption spectra of the C-dots were measured using a UV–vis spectrometer, with pure chloroform serving as the baseline.Three itraviolet absorption of C-dots were recorded under excitation wavelengths of 200–300nm, 350–450nm, and 550–750nm. Images of the C-dots were acquired using a transmission electron microscope(TEM) operating at an accelerating voltage of 200 kV.

### C-Dots Uptake Assay

Hybrid cells were seeded into 24-well and cultured overnight. After incubating the C-dots for 24 hours, the concentration-dependent cellular uptake was monitored using flow cytometry(excitation wavelength was 420nm, and the maximum emission wavelength was 680nm).

### CCK-8

A Cell Counting Kit-8 (CCK-8) assay (Beyotime,C0037) was employed to assess the impact of varying concentrations of carbon dots on the viability of hybrids. The cells were seeded into 96-well plates, with 100 μL of cell suspension added to each well. After adherence, the cells were treated with different dosages of carbon dots for 24h. Then the hybrids were incubated with the CCK-8 staining buffer for 1h. The optical density (OD) value was subsequently measured at 450nm.

### qRT-PCR

Total RNA was isolated using Trizol (Vazyme, R411) extraction. cDNA was synthesized from 1 μg of total RNA using HiScript Ⅱ Q RT SuperMix for qPCR (Vazyme, R123-01). qPCR was performed with ChamQ SYBR qPCR Master Mix (Vazyme, Q341-02) under the following conditions: an initial denaturation step at 95°C for 10 min, followed by 40 cycles of 95°C for 15s, 60°C for 30s, and 72°C for 30 s. Relative gene expression was calculated using the Ct method and normalized to GAPDH. The primer sequences utilized are provided in Table 1.

**Table 1.**
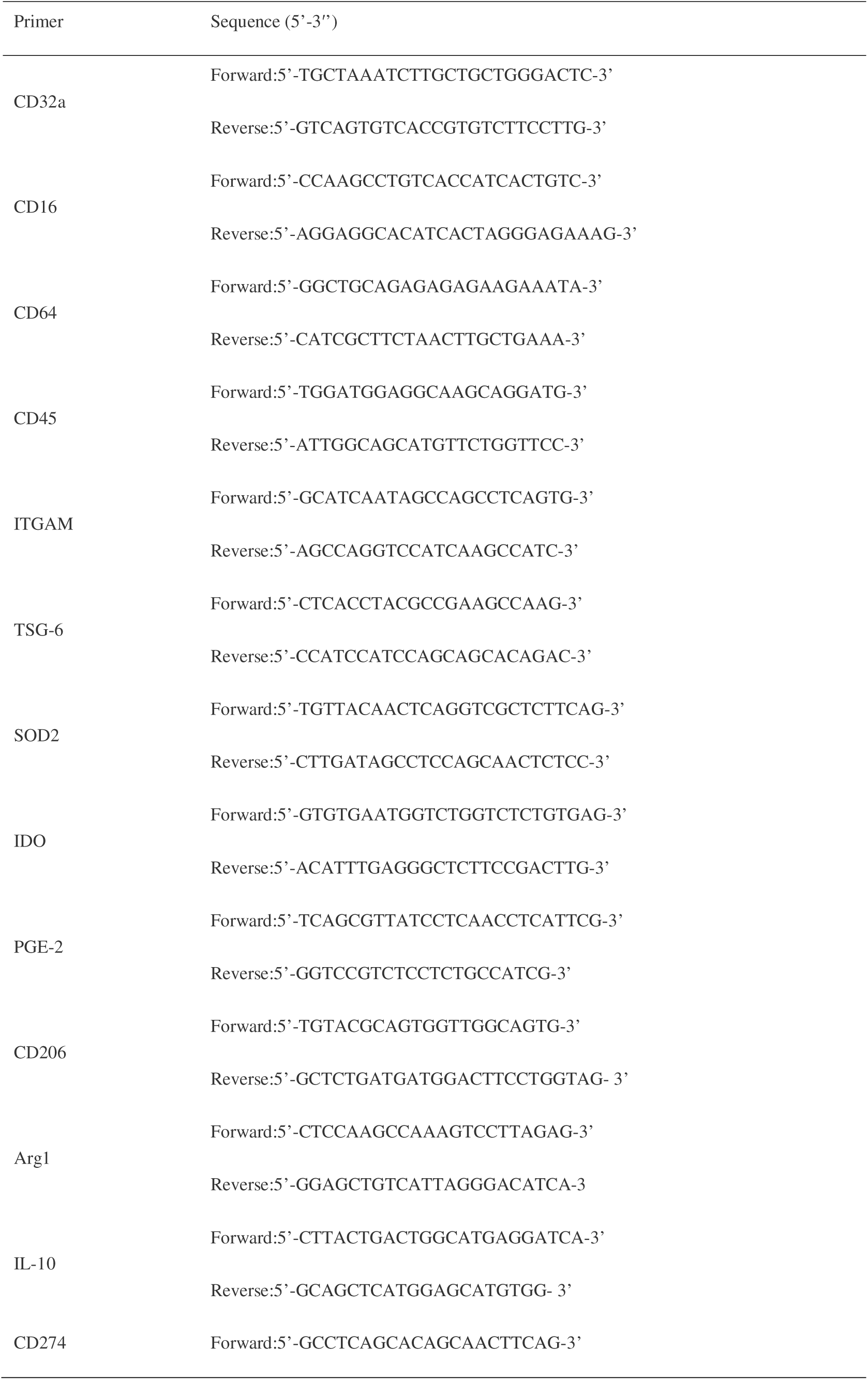
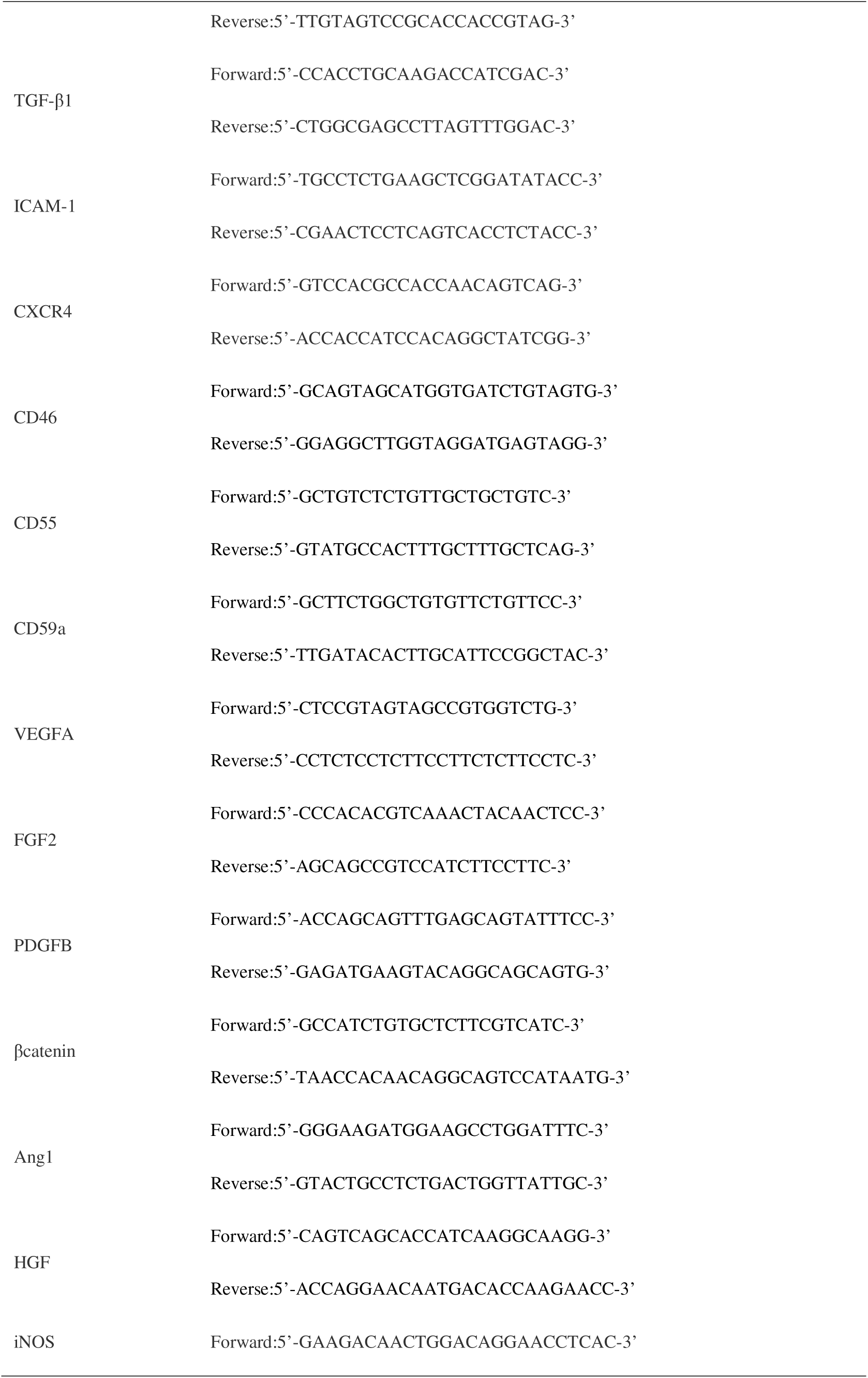
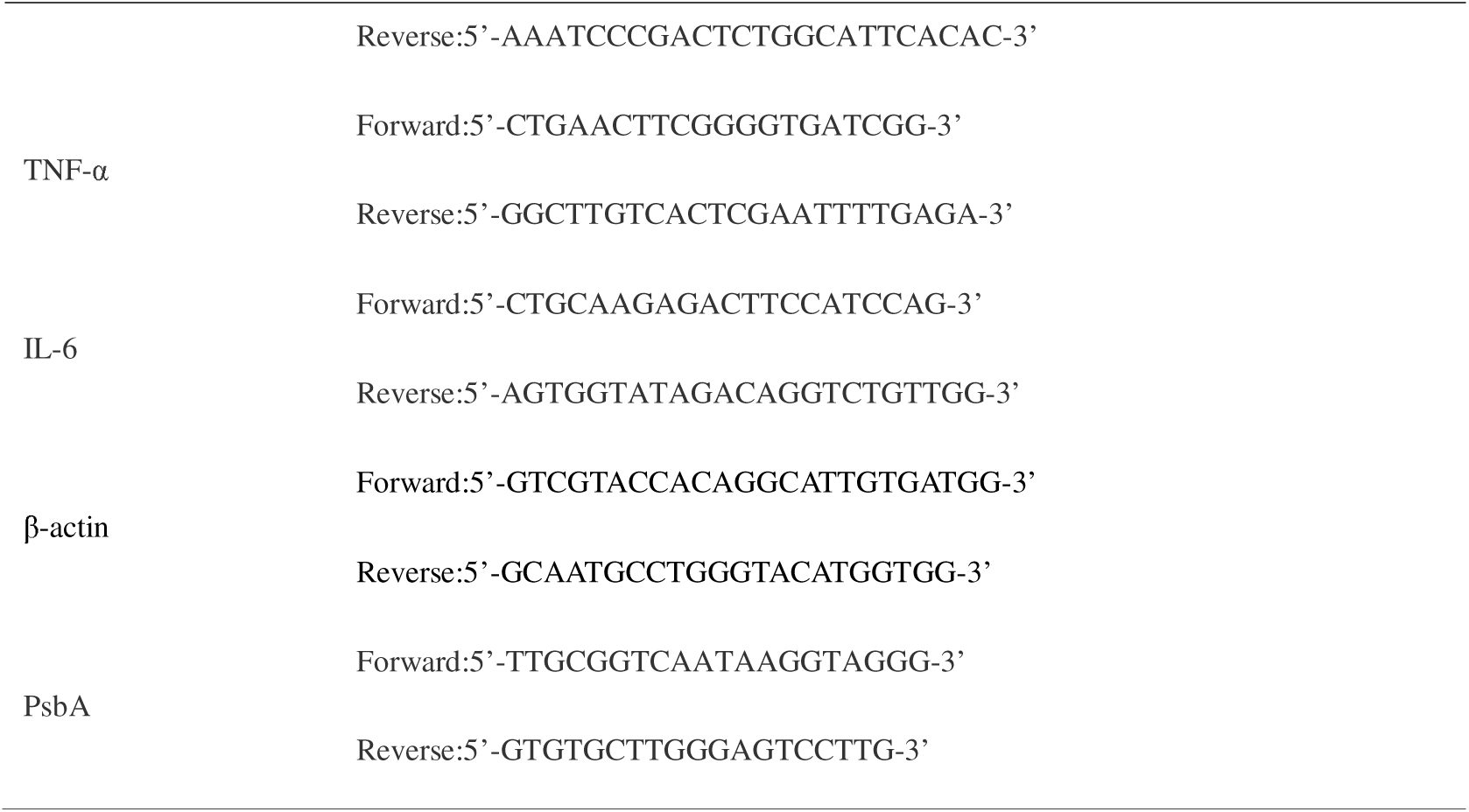
The primer sequences used for qRT-PCR.

### Statistical Analysis

Data are presented as mean ± standard deviation (SD) values. All experiments were conducted in triplicate. Statistical significance between the two groups was assessed using a two-tailed t-test, while multiple comparisons were evaluated using one-way analysis of variance (ANOVA). A p-value of less than 0.05 was deemed statistically significant. Statistical analyses were carried out using GraphPad version 8.0.

## Results

### Hybrids can be efficiently produced through BMSC and RAW264.7 fusion

To investigate whether BMSC can fuse with RAW264.7 cells through hACE2 and COVID-19-S, we first mixed PKH67-labeled BMSCs and PKH26-labeled RAW264.7. We observed double-positive cells, confirming the fusion process in the absence of apparent cell-in-cell or endocytosis structures as previously reported^21,22^ (Figure 1A). However, the hybrid was predominantly absent in the natural mixture without transfection treatment(data not show). It is noteworthy that RAW264.7 cells, when treated solely with the transfection reagent, were unable to reattach(Figure 1B). The fusion of BMSC and RAW264.7 cytoplasm was subsequently assessed using mitochondrial trackers. As anticipated, mitochondria from both cell types were observed simultaneously, indicating co-localization (Figure 1A). It is evident that the cell membrane and cytoplasm of the parents in the hybrid are completely fused and exist in a fluid state. Additionally, we analyzed the variations in the relative abundance of mitochondria in hybrid cells and discovered that the quantity of hybrid mitochondria was nearly three times higher compared to BMSC cells (Figure 1C).

**Figure 1.**
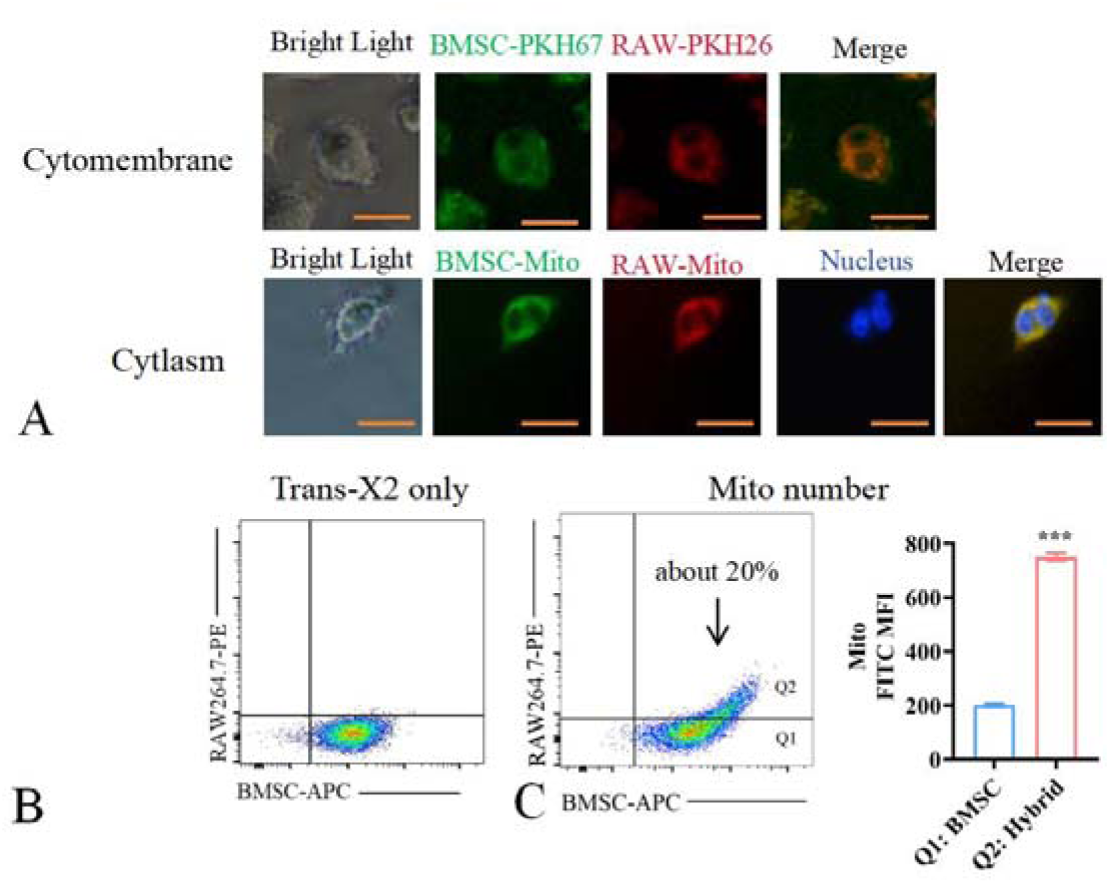
Mixed BMSCs and RAW264.7 form hybrids in vitro. A representative confocal micrograph illustrating the hybrids, showcasing the cytomembrane and cytoplasm of both BMSCs and RAW264.7(A). Flow cytometry analysis of BMSCs-DiD(APC)^+^RAW264.7(PKH26-PE)^+^ population after suffered from transfection reagent only(B). Flow cytometry plot to identify hybrid events and quantify mitochondrial statistics(C). Data are represented as means±SD (n=3), ***P<0.001 vs BMSC(two-tailed t-test). Scale bar 25μm.

While we detected roughly 20% of double-positive cells and a notable fraction of BMSCs via flow cytometry analysis (Figure 1C), we enriched these hybrids through magnetic sorting to facilitate a deeper examination of their functions. Superparamagnetic nanoparticles are readily taken up and can be manipulated via magnetic guidance. Previous research has shown that enhancing BMSCs and macrophages with magnetic nanoparticles significantly boosts their anti-inflammatory, reparative, and antioxidant properties ^23^. Importantly, these two cell types demonstrate different rates of nanoparticle uptake^24,25^. To evaluate cytotoxicity, various concentrations of Fe_3_O_4_ nanoparticles were co-cultured with the mixed cells for an initial duration of 24h. Our findings indicated that 15 μg/ml nanoparticles yielded optimal results, characterized by minimal levels of apoptosis(Figure 2A) and ROS(Figure 2B), stable MMP(Figure 2C), and satisfactory nanoparticle labeling as evidenced by Prussian blue staining (Figure 2D). Considering that hybrids may possess macrophage-like characteristics, we proposed that hybrids and BMSCs might demonstrate varying capacities for absorbing magnetic nanoparticles, which could influence their directional migration abilities in response to magnetic fields. We utilized this hypothesis by quickly directing the initial mixed cells to adhere to the wall, consequently removing unattached cells promptly. The purity of the hybrids was about 90%, as determined by flow cytometry analysis of the double staining rates using DiD and DiO (Figure 2E).

**Figure 2.**
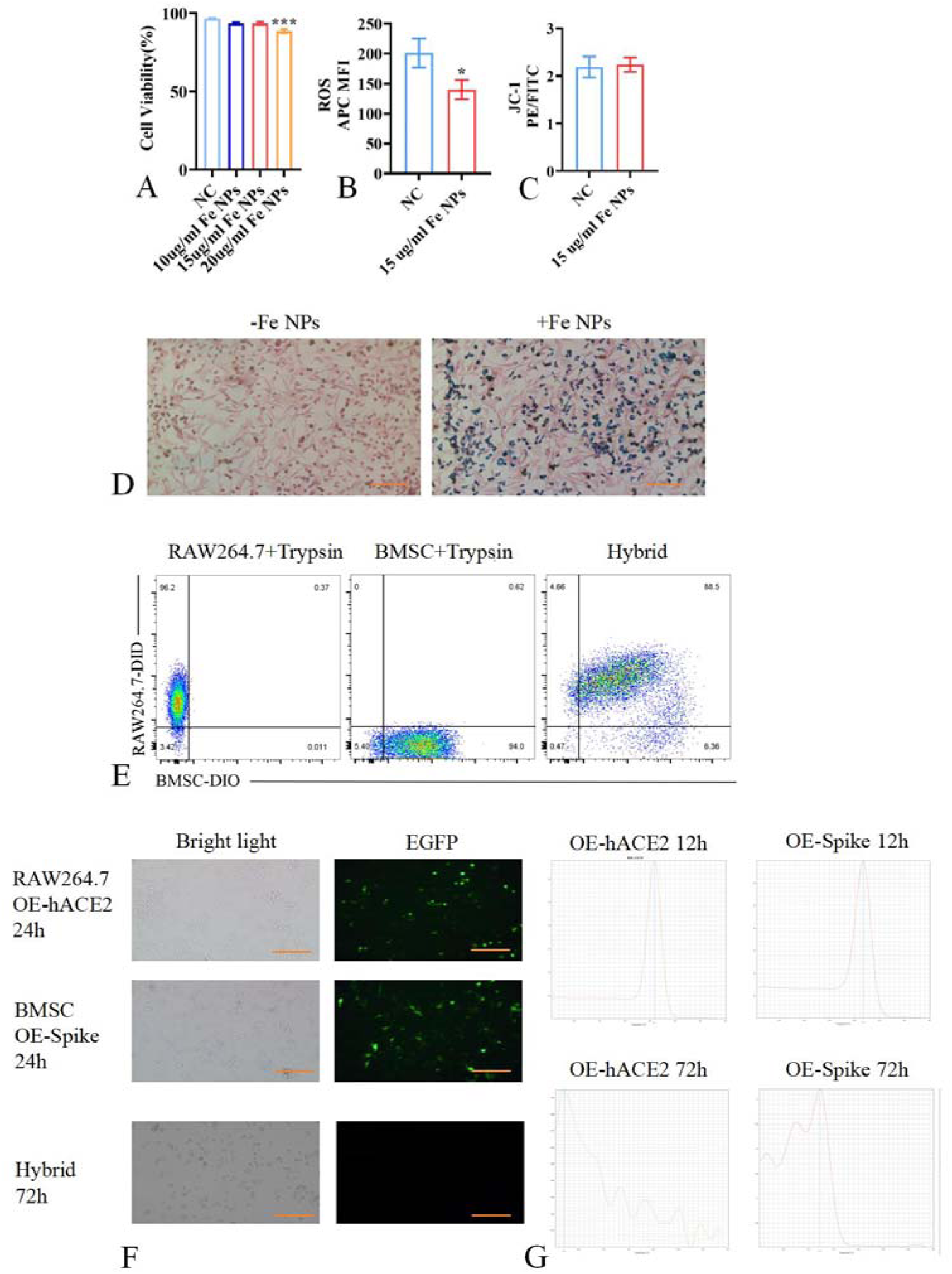
Hybrid purification and purification efficiency assessment. Flow cytometry detection of cytotoxicity(A) of Fe_3_O_4_ nanoparticles and their effects on ROS(B) and mitochondrial membrane potential(C). Determination of nanoparticle uptake rate through prussian blue staining(D). Flow cytometry measurement of DiD^+^DiO^+^ ratio(E). Immunofluorescence(F) and qRT-PCR(G) analysis of hACE2 and Spike expression. Data are represented as means±SD (n=3), *P<0.05, ***P<0.001 vs NC(two-tailed t-test). Scale bar 200μm.

In order to prevent any interference caused by hACE2 and Cov19-S, pcDNA3.1-CoV19S-EGFP and pcDNA3.1-hACE2-EGFP were separately introduced into BMSCs and RAW264.7 cells. The hybrids were then subjected to fusion and purification processes, after which they were cultured for 72h and analyzed via fluorescence microscopy and qRT-PCR. The lack of green fluorescence indicated that these linker proteins were not detectable at the protein level, while qRT-PCR verified their absence at the mRNA level(Figure 2F and 2G). These results suggest that the linker proteins do not affect subsequent functional assays following a culture period of 72h.

### Functional characterization of hybrids

To gain a deeper understanding of the characteristics of hybrids, scanning electron microscopy was utilized to investigate their morphological features. The findings suggested that the hybrids bore resemblance to RAW264.7 cells regarding their oval shape; however, a significant distinction was noted in their surface properties, with the hybrids displaying a rougher surface along with more protrusions and elongated structures. In comparison, BMSCs exhibited a flattened morphology reminiscent of fibroblasts(Figure 3A). Moreover, phalloidin staining results indicated that the distribution of the cytoskeleton in the hybrids closely mirrored that of RAW264.7 cells(Figure 3B). Subsequently, a colony formation assay was executed. BMSCs, RAW264.7 cells, and hybrids were plated in 6-well dishes. After a 7 day incubation period, crystal violet staining was performed. The results indicated that both hybrids and RAW264.7 cells successfully formed distinct colonies, though the hybrids’ colonies were somewhat smaller than those produced by RAW264.7 cells. Notably, BMSCs did not manage to generate visible colonies under the same experimental conditions(Figure 3C). To further clarify the functional differences between the three cells, a series of experiments was carried out. Initially, the capacity for glucose uptake was assessed. Hybrids had a significantly greater 2-NBDG uptake rate in comparison to BMSCs(Figure 3D). While the difference did not result in a two-fold increase, it still highlighted a clear functional divergence in glucose metabolism. Moreover, hybrids exhibited a notably greater phagocytic ability for erythrocyte fragments when compared to BMSCs and RAW264.7(Figure 3E). This outcome clearly underscored the clonogenic potential of the hybrids.

**Figure 3.**
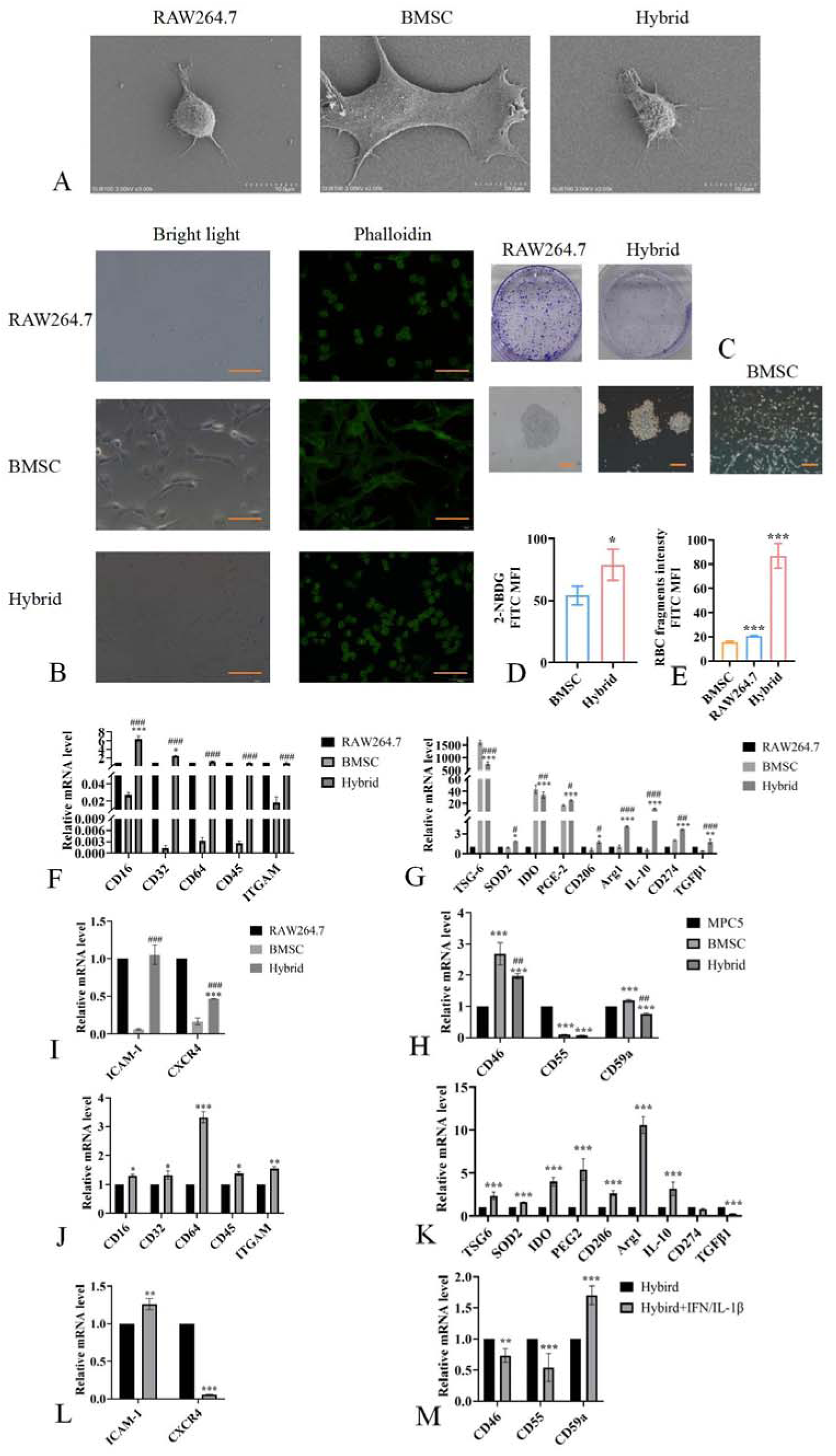
Characteristic analysis of hybrids. TEM images(A), phalloidin staining(B) and clonogenic assay(C) of RAW264.7, BMSC and hybrid. Flow cytometry detection of glucose uptake rate(D) and RBC fragment phagocytosis rate(E). qRT-PCR analysis of immunoregulatory factors(F-M). *P<0.05, ***P<0.001 vs BMSC in D and E(two-tailed t-test); *P<0.05, **P<0.01, ***P<0.001 vs RAW264.7, ^#^P<0.05, ^##^P<0.01, ^###^P<0.001 vs BMSC in F-H; ***P<0.001 vs MPC5, ^##^P<0.01vs BMSC in I; *P<0.05, **P<0.01, ***P<0.001 vs hybrid in J-M(One-way ANOVA Tukey’s multiple comparison). Scale bar 200μm.

To evaluate whether the hybrids preserved the anticipated functions of both parental cell types, a qPCR analysis was performed. In terms of FcγRs, the hybrids showed a notable upregulation of CD16 and CD32 when compared to RAW264.7 cells. In comparison to BMSCs, the hybrids exhibited an upregulation of CD16, CD32, CD64, CD45, and ITGAM (Figure 3F). As for anti-inflammatory and antioxidant factors, relative to RAW264.7 cells, the hybrids displayed increased levels of TSG-6, SOD2, IDO, PGE-2, CD206, Arg1, IL-10, CD274, and TGFβ1. When compared with BMSCs, the hybrids likewise showed an increase in SOD2, PGE-2, CD206, Arg1, IL-10, CD274, and TGFβ1, whereas TSG-6 and IDO were found to be downregulated (Figure 3G). In relation to adhesion factors, the hybrids exhibited elevated ICAM-1 levels compared to both RAW264.7 cells and BMSCs; however, CXCR4 levels were significantly reduced in comparison with BMSCs (Figure 3H). In relation to complement inhibitors, the hybrids showed elevated CD46 levels, but diminished quantities of CD55 and CD59a when evaluated against MPC5 cells, alongside lowered levels of both CD46 and CD59a in comparison to BMSCs (Figure 3I).

Following the IL-1β/IFNγ stimulation of hybrids, there was a notable upregulation of CD16, CD32, CD64, CD45, and ITGAM, (Figure 3J). Heightened levels of TSG-6, SOD2, IDO, PGE-2, CD206, Arg1, and IL-10, while TGFβ1 were found to be lower (Figure 3K). With respect to adhesion factors, a slight increase was seen in ICAM-1, whereas CXCR4 exhibited a significant decrease (Figure 3L). As for complement inhibitors, CD59a showed an increase, while CD46 and CD55 demonstrated decreases (Figure 3M). Collectively, these results robustly depict that the hybrids possess a unique array of functions, showcasing enhanced anti-inflammatory and antioxidant abilities, especially in the context of inflammatory stimulation.

### Hybrids clear IgG along with promoting injury repair

To investigate the clearance effect of hybrids on IgG complexes, we initially evaluated their capacity to internalize free IgG-FITC using immunofluorescence detection. Our findings showed that the green fluorescence was concentrated in the cytoplasm of the hybrids, indicating that they can efficiently endocytose IgG(Figure 4A). Additionally, flow cytometry analysis demonstrated that the uptake of IgG-FITC by the hybrids was markedly increased after pre-activation with IL-1β/IFNγ(Figure 4B). Compared to the group that is non-phagocytic, the phagocytic group with IgG exhibited notably higher expression of CD16, CD64, ITGAM, TSG-6, SOD2, IDO, PGE-2, CD206, Arg1, IL-10,

**Figure 4.**
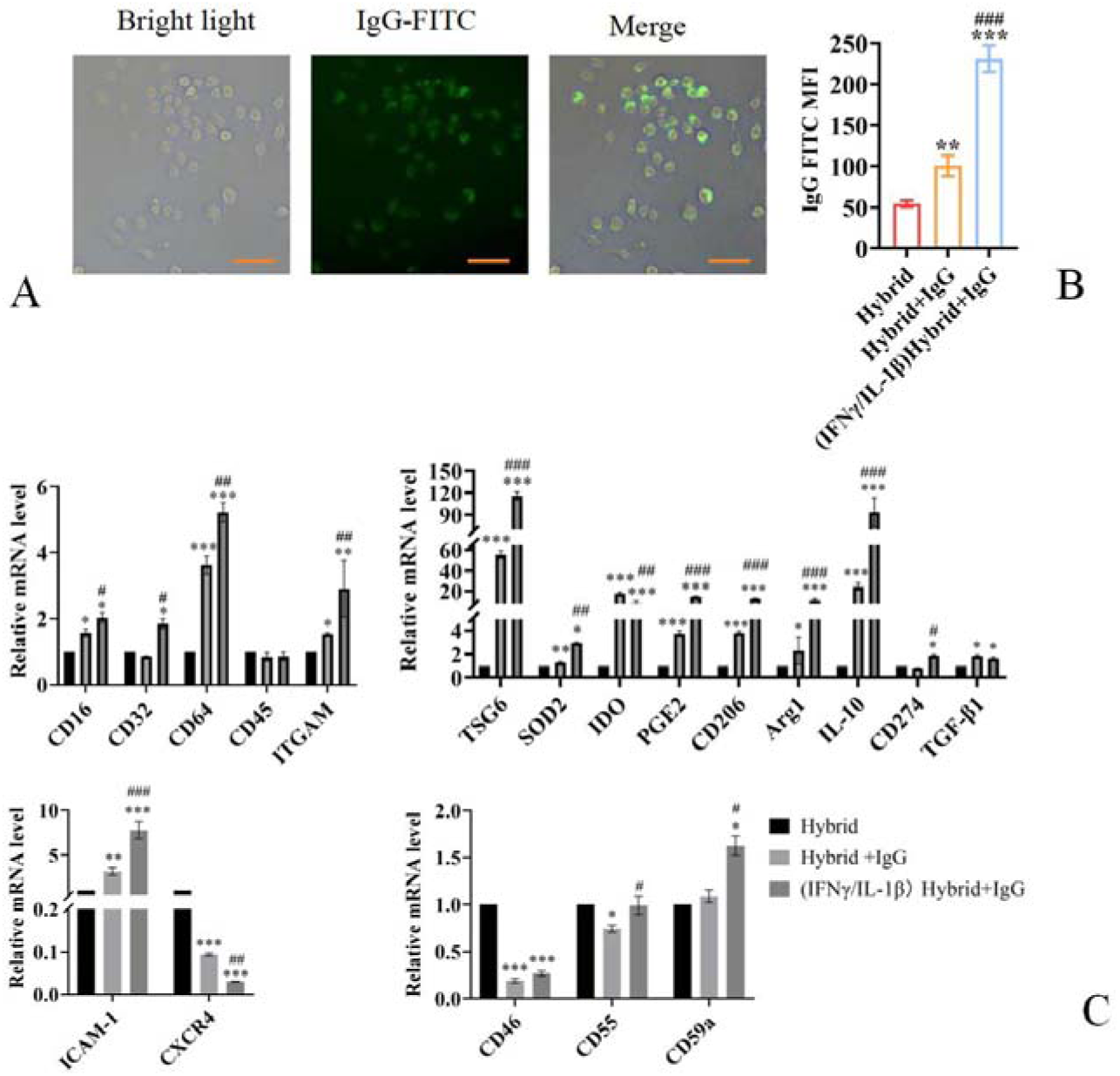
Hybrids clear free IgG. Immunofluorescence and qRT-PCR analysis of IgG-FITC distribution(A) in hybrids and the expression of immunoregulatory factors(B). Data are represented as means±SD (n=3), *P<0.05, **P<0.01, ***P<0.001 vs hybrid, ^#^P<0.05, ^##^P<0.01, ^###^P<0.001 vs hybrid+IgG in B and C(One-way ANOVA Tukey’s multiple comparison). Scale bar 100μm.

TGFβ1, ICAM-1, and CD59a, in contrast, there was a significant reduction in the levels of CXCR4, CD46, and CD55. Additionally, when pre-activated with IL-1β/IFNγ, there was further marked increase in the expression of CD16, CD32, CD64, ITGAM, TSG-6, SOD2, PGE-2, CD206, Arg1, IL-10, CD274, ICAM-1, CD46, CD55, and CD59a, whereas IDO, TGFβ1, and CXCR4 experienced an even greater decline(Figure 4C).

Following this, we developed a model of IgG deposition to study the effects of hybrids on kidney organoids exposed to inflammatory injury. In this research, we focused on podocytes due to their critical role in various immune-mediated glomerulopathies, including lupus glomerulonephritis and transplant glomerulopathy^26,27^. Our results showed that treating MPC5 cells with DOX led to a significant increase in the rate of apoptosis, a pronounced reduction in mitochondrial membrane potential, a significant uptick in ROS levels, and notable increases in the expression of IL-6, TNF-α, IL-1β, and IFNγ (Figure 5A-D). In line with previous studies^28^, flow cytometry analysis indicated that both the proportion of CD80-positive cells and their expression levels were significantly elevated (Figure 5E).

**Figure 5.**
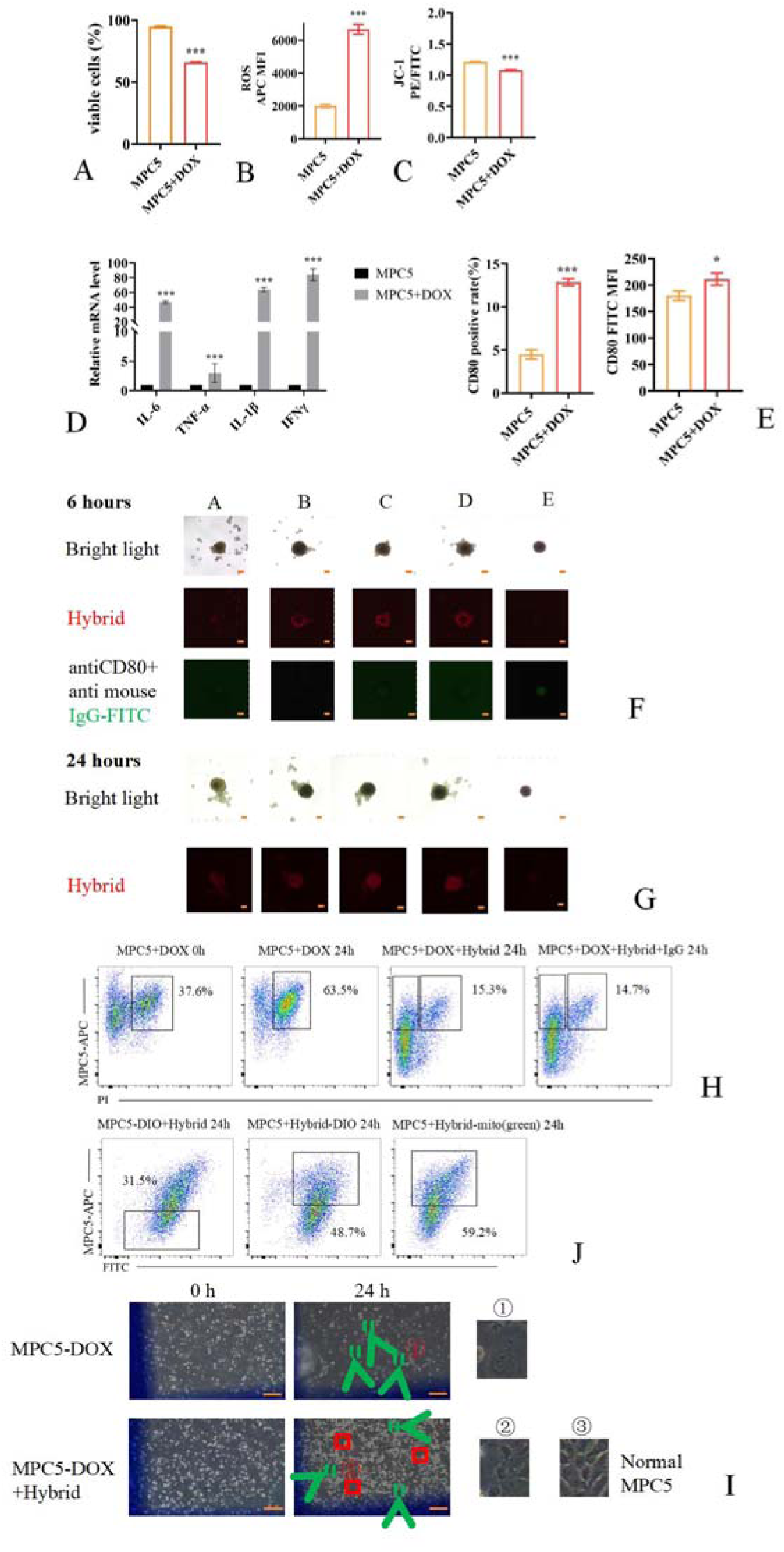
Hybrids infiltrate kidney organoids, eliminate deposited IgG and reverse MPC5 inflammatory damage. Flow cytometry analysis of apoptosis(A), ROS(B), JC-1(C), CD80^+^MPC5 population and CD80 expression(E). qRT-PCR detection of IL-6, TNF-α, IL-1β and IFN-γ(D). Confocal images of hybrids infiltration and IgG disappearance(F and G). Flow cytometry analysis of MPC5 viability under co-culturing with hybrids(H) and the mitochondrial and vesicle transport status between hybrids and MPC5 cells(J). Images of changes in MPC5 morphology and quantity(H). Data are represented as means±SD (n=3). *P<0.05, **P<0.01, ***P<0.001 vs MPC5(two-tailed t-test). Scale bar 200μm in I, 100μm in F and G.

Based on these results, we proceeded to label DOX-treated MPC5 cells with a rabbit anti-mouse CD80 primary antibody, followed by a mouse anti-rabbit IgG-488 fluorescent secondary antibody. We then created 3D microspheres using nucleic acid scaffolds. Hybrids marked with Cell Deep Red tracker were co-cultured with these microspheres. It was evident that the hybrids demonstrated enhanced infiltration towards the DOX+IgG deposition group, successfully clearing IgG, with green fluorescence absentwithin 6 h (Figure 5F). Additionally, they exhibited increased invasiveness after 24h (Figure 5G). Notably, pre-treatment with IFNγ/IL-1β did not yield any observable effects.

To further clarify the effects of hybrids on DOX-treated MPC5 cells, we labeled MPC5 with the Cell Deep Red tracker and co-cultured it with hybrids. It was observed that the apoptosis rate of MPC5 cells treated with DOX continued to increase after 24 h of culture, whereas the apoptosis rate in the co-culture group significantly decreased. Pre-labeling with the IgG antibody had no effect on the viability of MPC5 cells (Figure 5H). These results indicate that in the presence of the IgG antibody, the hybrids did not induce ADCC effect on the inflammatory-damaged MPC5 cells, while simultaneously improving cell viability.

Besides, the DOX-treated MPC5 group displayed swelling and enlargement of the nucleus. Conversely, the co-culture group containing hybrids exhibited a notable increase in intercellular space, and some MPC5 cells appeared to revert towards their typical morphology (Figure 5I). Integrating the findings from the previous section suggests that hybrids may phagocytize the more severely impaired MPC5 cells while simultaneously repairing those that are less affected.

Further testing revealed that DOX-treated MPC5 cells significantly incorporated the hybrid membrane components as well as their mitochondrial(Figure 5J). Similarly, the cell membranes of DOX-treated MPC5 cells were extensively taken up by hybrid cells. These results suggest that the hybrids may influence MPC5 cells through vesicle/mitochondrial transfer and efferocytosis.

### Hybrids exhibit efferocytosis capability

Macrophages play a critical role in the timely resolution of inflammation and the enhancement of anti-inflammatory effects by engulfing apoptotic cells through efferocytosis, thereby contributing to the immune regulation of tissue repair^29^. To further investigate whether hybrids exhibit a similar efferocytic function as macrophages, we co-cultured MPC5-ACs with hybrids. Furthermore, the proliferative ability of hybrids that had engulfed ACs was significantly reduced (Figure 6C). qRT-PCR analysis revealed that, compared to the non-ACs group, the levels of SOD2, Arg1, IL-10, CD274, TGF-β1, VEGFA, FGF2, PDGFB and βcatenin were significantly elevated in the phagocytosis group, while the levels of IDO and PGE2 exhibited a slight decrease (Figure 6D).

**Figure 6.**
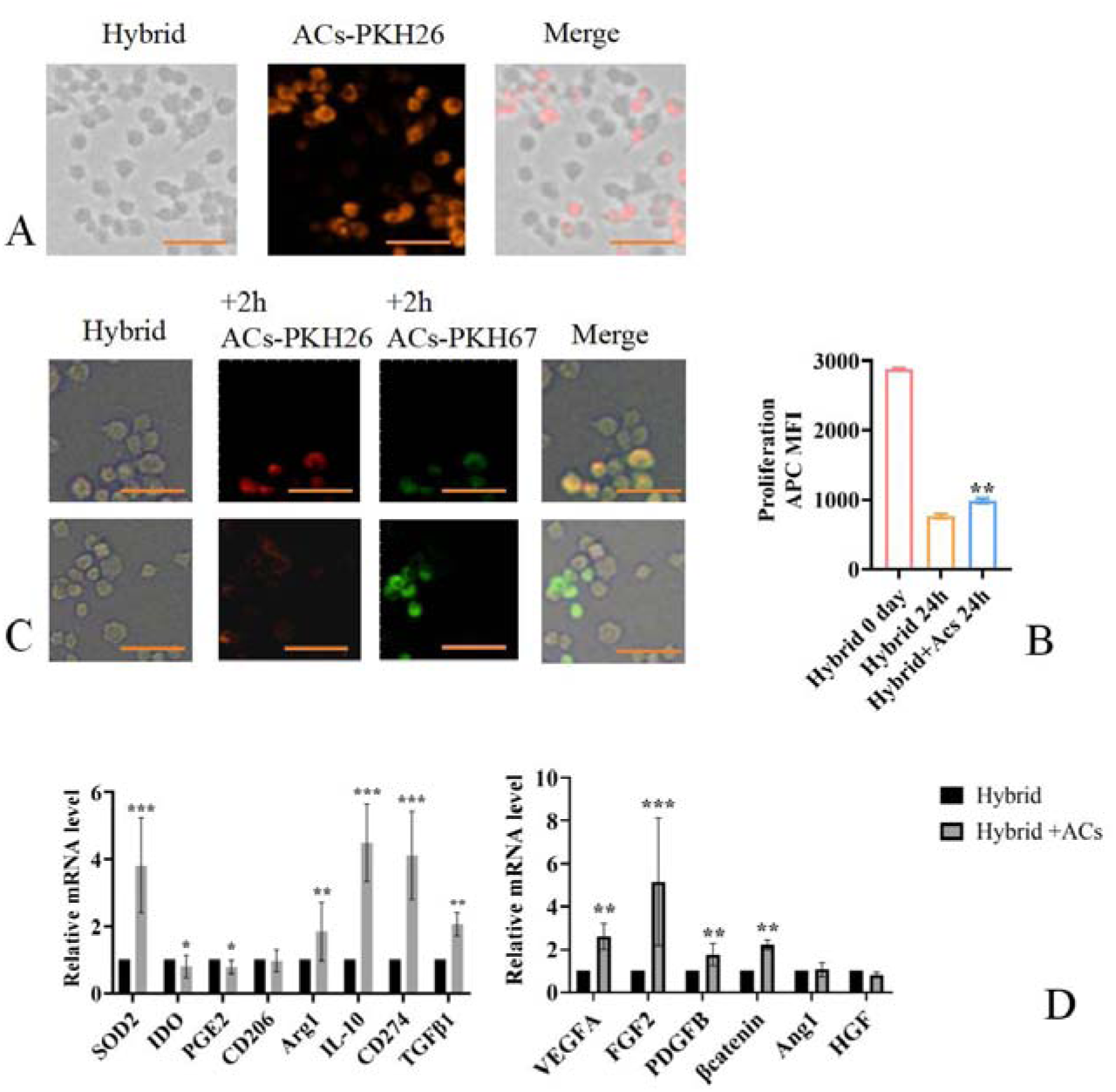
Hybrids efferocytosis. Confocal images of the phagocytic effect of hybrids on MPC5-ACs(A). Flow cytometry analysis of the changes in hybrids proliferation(B). Confocal images of the secondary phagocytic effect of hybrids on MPC5-ACs(C). qRT-PCR detection of anti-inflammatory and pro-repair factors(D). Data are represented as means±SD (n=3). *P<0.05, **P<0.01, ***P<0.001 vs hybrids(two-tailed t-test). Scale bar 100μm.

### Hybrids enhance the immunosuppressive phenotype of iDCs

Between the peritubular capillaries and epithelial cells, there exists a significant distribution of dendritic cells, which can directly interact with epithelial cells, endothelial cells, and circulating immune effector cells. Based on functional differences, DCs can be categorized into pro-inflammatory (mDCs) and anti-inflammatory (iDCs) types and the dysregulation of DCs is a critical mechanism in the onset and progression of immune-related kidney diseases^30^. MSCs enhance nitric oxide synthase activity by stimulating renal iDCs to secrete IL-10, thereby exerting a protective effect on the kidneys. MSCs not only inhibit the maturation process of DCs but also promote their transformation into an immunosuppressive phenotype^29^. Our study aims to investigate whether hybrids inherit the functional characteristics of MSCs and to explore their potential impact on iDCs.

The findings from our study revealed a marked reduction in the proliferation of iDCs (Figure 7A), which was accompanied by a significant increase in PDL-1 level (Figure 7B) after co-culturing with hybrids. Additionally, these effects became even more evident following prior stimulation with IFN-γ/IL-1β.

**Figure 7.**
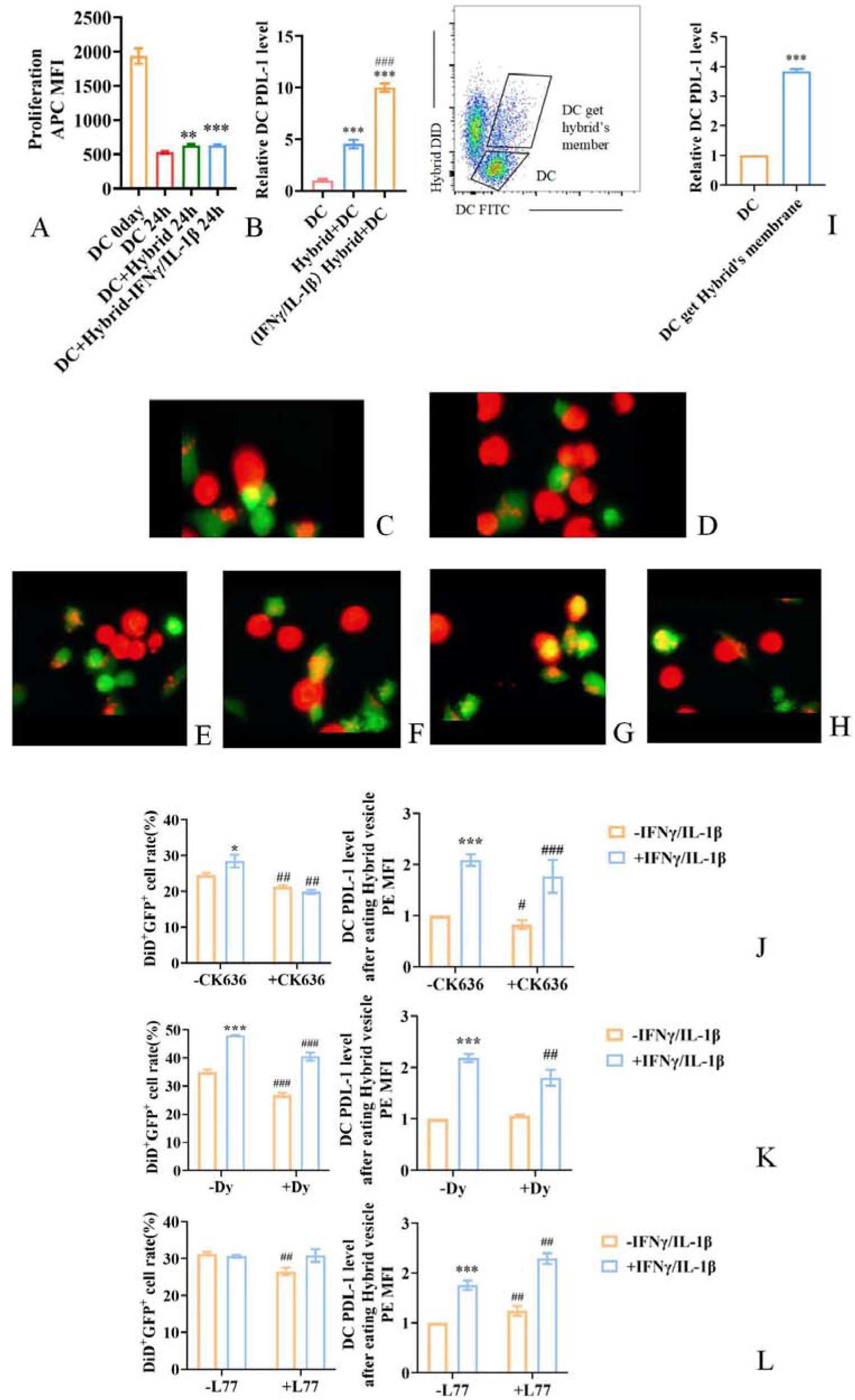
Hybrids enhance the immunosuppressive function of iDCs and their underlying mechanisms. Flow cytometry sorting and detection of proliferation(A) and PDL-1 levels in total iDCs(B) and the iDCs with obtaining hybrid membrane components(I) after co-culture with hybrids. Live-cell imaging observations revealed that hybrids influence iDCs through cell migration(C), nanotube interactions(D and G), efferocytosis(E), label tracking(F), and label pulling(H). The impact of CK636(J), Dynasore(K) and L778123(L) on the proportion of iDCs cells with hybrid membrane components (DiD^+^GFP^+^) and PDL-1 levels. Data are represented as means±SD (n=3). **P<0.01, ***P<0.001 vs DC 24h in A(two-tailed t-test); ***P<0.001 vs DC and ^###^P<0.001 vs Hybrid+DC in B(One-way ANOVA Tukey’s multiple comparison); ***P<0.001 vs DC in I(two-tailed t-test); *P<0.05, **P<0.01 vs -IFN-γ/IL-1β and ^#^P<0.05, ^##^P<0.01, ^###^P<0.001 vs -CK636/-Dy/-L77 in J-L(two-tailed t-test).

Live-cell imaging was conducted to study the dynamic cellular behaviour between hybrids and iDCs. The data indicated that hybrids produced a significant number of vesicles during their migration, which were later absorbed by iDCs(Figure 7C). In addition, the hybrids demonstrated an ability to transfer vesicles to iDCs through nanotubular structures (Figure 7D). Moreover, hybrids prompted iDCs to break apart into various small vesicles, which were then taken up (Figure 7E). Hybrids actively targeted iDCs that had consumed their membrane components (Figure 7F), and the membrane elements of hybrids could be exchanged between iDCs through nanotubular connections (Figure 7G). Furthermore, hybrids applied a pulling force on iDCs that had ingested their membrane components, thus obstructing their migration (Figure 7H). These results imply that hybrids may engage with iDCs through migrasomes, nanotubes, efferocytosis, and cellular traction. Alongside the results shown in Figure 7B, we hypothesize that hybrids might affect PDL-1 expression in iDCs by delivering vesicles through migrasomes and nanotubes.

To validate the above hypothesis, we co-cultured DID-labeled hybrids with iDCs and observed that the PDL-1 level in iDCs which acquired hybrid vesicles was significantly higher than that in the non-acquired group (Figure 7I). To further elucidate the pathways influencing PDL-1 expression in iDCs, we employed L-778123, Dynasore, and CK636 in our study. All three drugs were found to inhibit the uptake of hybrid vesicles by iDCs. Notably, after pre-treatment with L-778123, the PDL-1 level in iDCs that internalized hybrid vesicles was increased. In contrast, pre-treatment with Dynasore did not result in a significant difference in PDL-1 level in iDCs that internalized hybrid vesicles. Additionally, following pre-treatment with CK636, the PDL-1 level in iDCs that took up hybrid vesicles significantly decreased. Furthermore, pre-treatment of hybrids with IFNγ/IL-1β can make the effects of these three inhibitors on the expression of PDL-1 in DC cells more significant.(Figure 7J-L). These findings suggest that hybrids primarily enhance the upregulation of PDL-1 expression in iDCs through the migrasome pathway.

### Hybrids maintain anti-inflammatory capacity under LPS/IFNγ

Macrophages can undergo polarization towards the M1 phenotype when exposed to inflammatory stimuli. Our objective was to assess whether hybrids exhibit this characteristic by administering LPS/IFNγ; however, we did not observe any significant morphological alterations typical of M1-type macrophages (Figure 8A). Furthermore, the LPS/IFN-γ-treated hybrids exhibited a significant increase in the levels of CD206, Arg1, and IL-10, while showing a decreased expression of TNF-α and IL-6 compared to LPS/IFN-γ-treated RAW264.7 cells. (Figure 8B). These results indicate that hybrids are capable of maintaining a distinct anti-inflammatory profile during LPS/IFNγ exposure.

**Figure 8.**
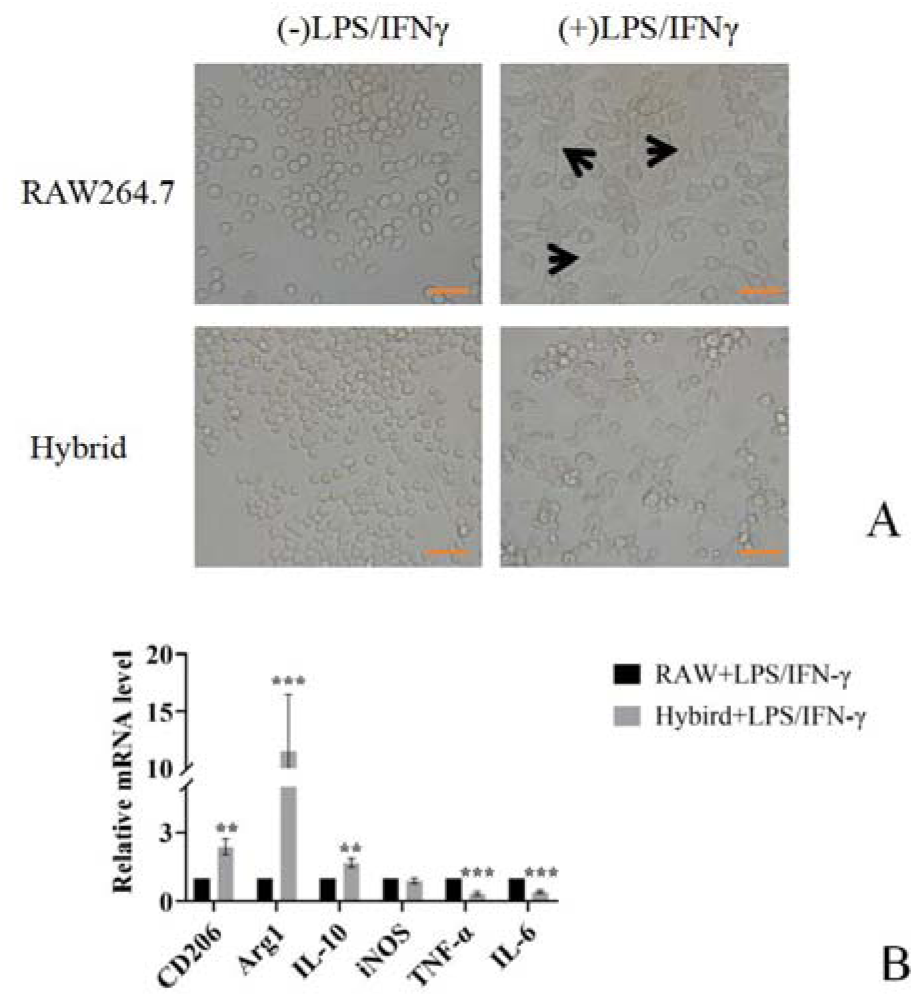
Hybrids still possess antioxidant capacity under LPS/IFNγ exposure. Microscopic images of cellular morphological changes(A). qRT-PCR analysis of CD206, Arg1, IL-10, iNOS, TNF-α and IL-6(B). Data are represented as means±SD (n=3). **P<0.01, ***P<0.001 vs RAW+LPS/IFN-γ(two-tailed t-test). Scale bar 100μm.

### Hybrids-derived vesicles can target IgG deposition areas and adsorb IL-6

In recent years, artificial vesicles derived from cells have attracted considerable interest across various domains, including immune regulation, inflammation management, tissue repair enhancement, and anti-tumor therapies. These vesicles are noted for their cost effectiveness in preparation, minimal immunogenic response, operational simplicity, and ease of engineering modifications, acting as an extension of cellular capabilities^31^. In our research, we integrated membranes sourced from BMSCs and macrophages into our hybrids. Following this, we utilized the artificial membrane extrusion technique to create vesicles derived from these hybrids, aiming to explore their role in shaping the inflammatory microenvironment. Observations made through transmission electron microscopy indicated that these vesicles displayed a well-defined nanostructural morphology (Figure 9A). Nanoparticle tracking analysis showed the particles’ size to be 197.7 ± 79.1 nm. In addition, Western blot assessments demonstrated that CD16 and CD64 expression levels in hybrid-derived nanovesicles were akin to those found in RAW264.7 nanovesicles, while PDL-1 levels were similar to that of BMSC-derived nanovesicles. Notably, the IL-6R level was significantly higher when compared to both RAW264.7 and BMSC-derived nanovesicles (Figure 9B). These findings imply that hybrid-derived nanovesicles exhibit elevated expression of CD16 and CD64, which can bind to the Fc region of IgG antibodies, alongside PDL-1 that possesses immunosuppressive characteristics, and IL-6R that interacts with IL-6.

**Figure 9.**
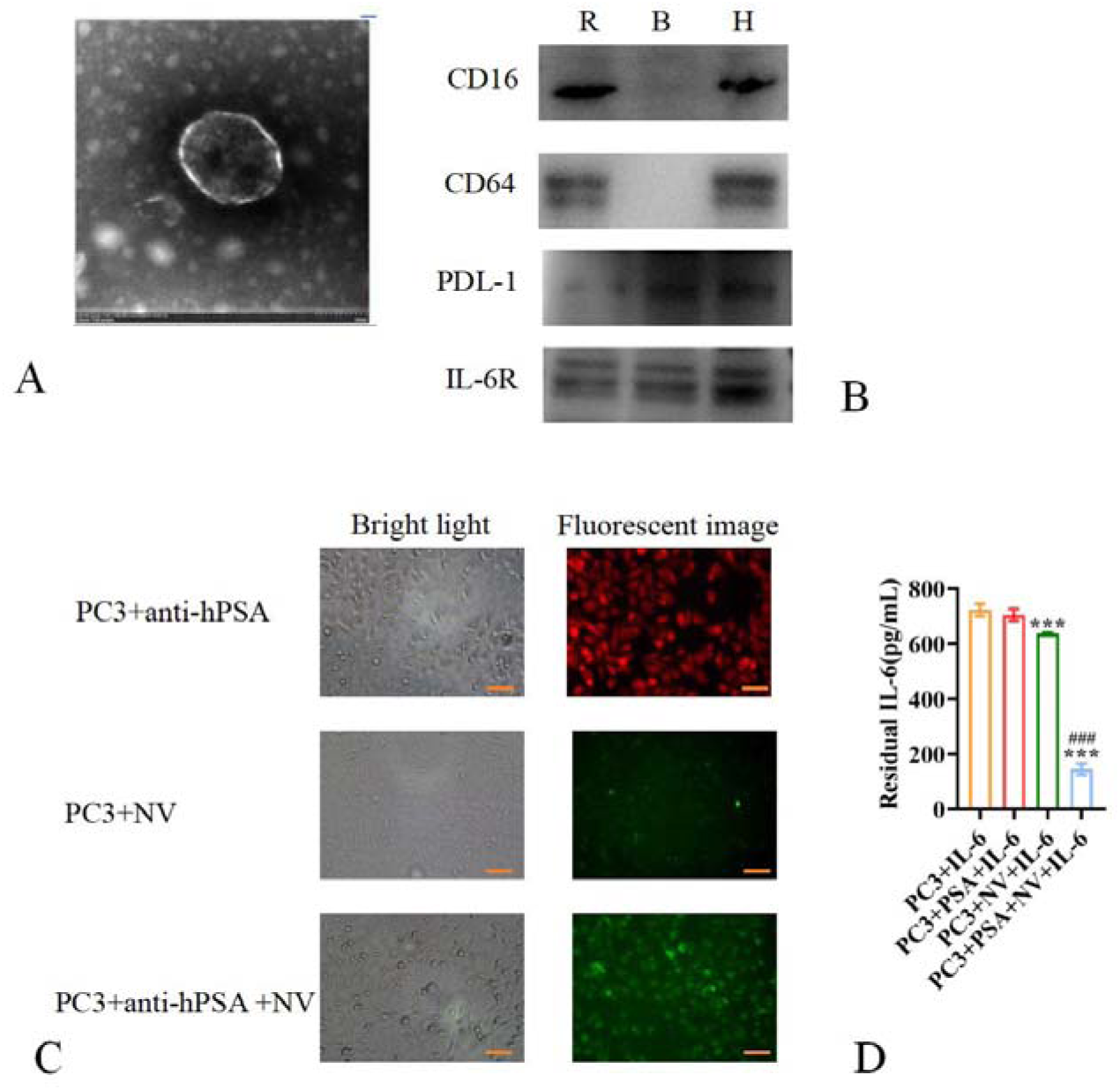
Hybrids-derived vesicles effectively adsorb free IL-6. TEM images of hybrids-derived vesicles(A), Western blot analysis of CD16, CD64, PDL-1 and IL-6R levels(B). Microscopic images of the targeting binding ability of hybrids-derived vesicles to IgG(C). ELISA analysis of IL-6 concentration(D). Data are represented as means±SD (n=3). ***P<0.001 vs PC3+IL-6, ^###^P<0.001 vs PC3+NV+IL-6(two-tailed t-test). Scale bar 50μm.

To assess this hypothesis, we initially employed mouse anti-human PSA antibodies to label fixed PC3 cells, aiming to reduce non-specific reactions and the effects of IL-6 produced by the cells themselves. Immunofluorescence analysis demonstrated that unlabeled PC3 cells did not efficiently interact with hybrid-NVs, while mouse anti-PSA antibody-labeled PC3 cells showed a markedly enhanced binding to hybrid-NVs. This observation confirms that hybrid-NVs do possess the capability to target IgG (Figure 9C). Following this, we co-cultured mouse IL-6 with fixed PC3 cells, organizing them into four distinct groups: ⍰PC3+IL-6; ⍰PC3+PSA+IL-6; ⍰PC3+NV+IL-6; and ⍰PC3+PSA+NV+IL-6. We utilized ELISA to assess the remaining IL-6 concentrations in the supernatant. Our findings indicated that the IL-6 levels in group ⍰ were significantly lower than those in group ⍰. These findings imply that hybrid-NVs directed at the IgG region can efficiently adsorb IL-6, thus mitigating its inflammatory damage to cells (Figure 9D).

### Hybrids loaded with chloroplasts promote wound repair under light-driven

To broaden the use of hybrids, our objective was to modify them. We initially targeted the area of wound care, where MSCs and their derivatives exhibit remarkable potential for transformative uses^32^. The effectiveness of MSC transplantation in facilitating the healing of both skin and internal wounds has been notably significant. However, a major obstacle in clinical settings is the limitation of the therapeutic efficacy duration of MSCs, given that the transplanted MSCs in the wound environment also experience hypoxia and oxidative stress^33,34^.

Studies have discovered that spinach thylakoids encapsulated within chondrocyte membranes (CM-NTUs) can elevate ATP and NADPH levels in chondrocytes when exposed to light, rectify energy discrepancies, and restore cellular metabolism, thereby preventing the pathological advancement of osteoarthritis^35^. Furthermore, experiments indicated that macrophages are capable of taking up CM-NTUs. Nonetheless, the extraction and preparation methods for CM-NTUs are intricate, and NTUs only retain partial chloroplast functions. Transplanting whole chloroplasts into mammalian cells for optimal function could offer greater advantages for clinical usage. Later, we co-incubated intact chloroplasts isolated from spinach with the hybrids. Dynamic imaging of live cells demonstrated that the hybrids actively internalized the intact chloroplasts (Figure 10A) within 3h, and components of the chloroplast membrane and ctDNA (PsbA) were remained detectable in the cells of the hybrids after 24h(Figure 10B-C). A further analysis of the pathway for chloroplast uptake indicated that pretreating the hybrid cells with the dynamin inhibitor dynasore (80 μg/ml) for 8h markedly decreased the internalization of chloroplasts (Figure 10D), suggesting that the uptake of chloroplasts by the hybrids relies on a dynamin-dependent mechanism.

**Figure 10.**
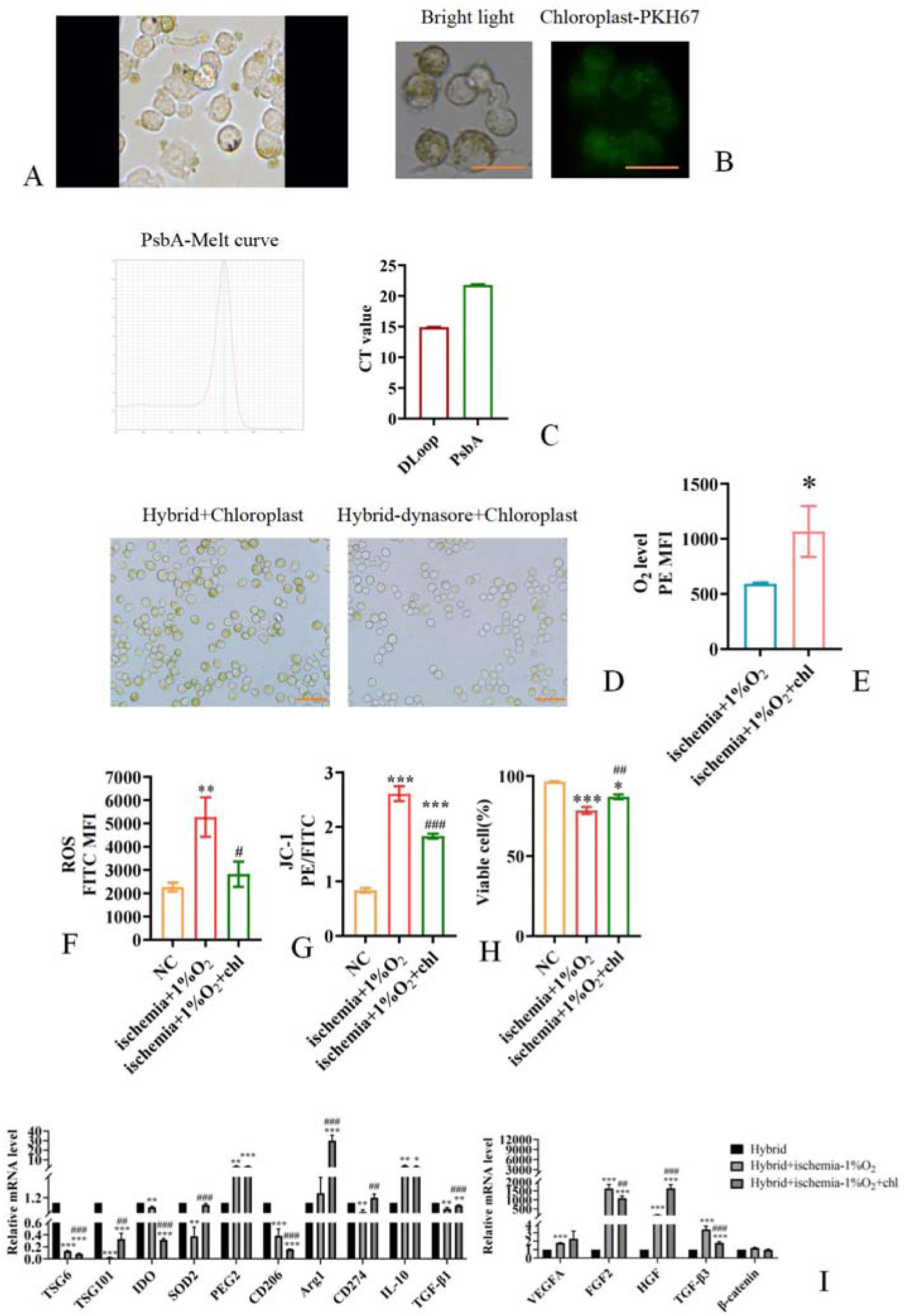
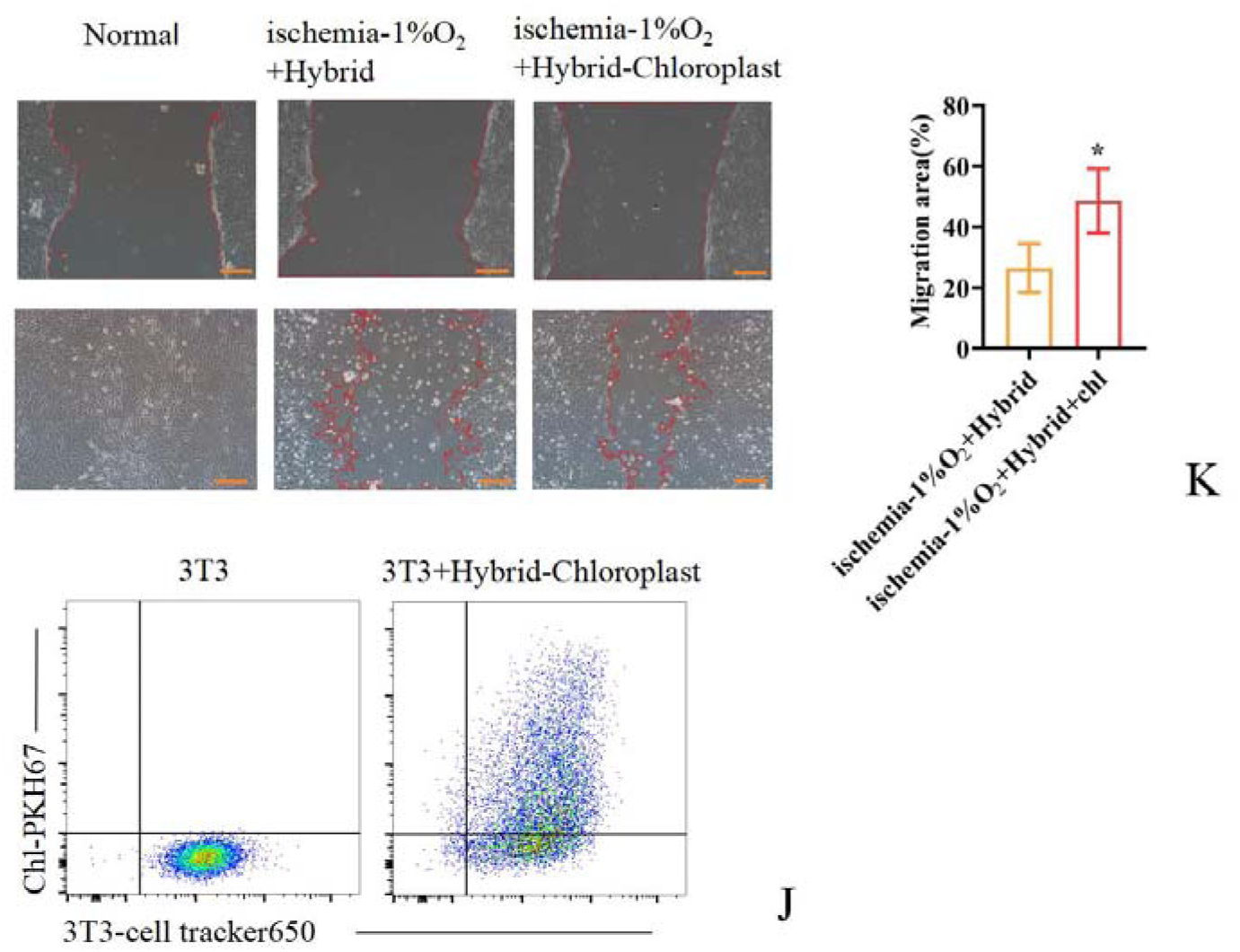
The chloroplasts carried by hybrids facilitate adaptation to the wound microenvironment. Live-cell imaging of the process of chloroplasts uptake by hybrids(A). Images of the localization of chloroplasts in hybrids(B). Melt curve and CT value of PsbA(C). Images of the variation in chloroplast content in hybrids(D). Flow cytometry analysis of intracellular oxygen content(E), ROS level(F), JC-1(G), apoptosis(H). qRT-PCR measurement of anti-inflammatory and pro-repair factors levels(I). Flow cytometry detection of chloroplast transfer between hybrids and 3T3 cells(J). Scratch assay and statistical plot for detecting changes in 3T3 cell migration(K). Data are represented as means±SD (n=3). *P<0.05 vs ischemia+1%O2 in E(two-tailed t-test); **P<0.05, ***P<0.001 vs NC, ^#^P<0.05, ^##^P<0.01, ^###^P<0.001 vs ischemia+1%O_2_ in F-H(One-way ANOVA Tukey’s multiple comparison). chl: Chloroplast. Scale bar 50μm in B, 100μm in D and 200μm in K.

Surprisingly, the hybrids loaded with chloroplasts demonstrated a significant increase in intracellular oxygen content, a reduction in ROS level, restoration of mitochondrial membrane potential, and a decrease in the rate of cell apoptosis under light treatment in a low-oxygen and starvation culture condition(Figure 10E-H). Additionally, the levels of TSG101, SOD2, Arg1, PDL-1, TGF-β1, and HGF were significantly elevated under chloroplasts loading(Figure 10I). Furthermore, through transwell chamber co-culture experiments, it was found that the hybrids could transfer the absorbed chloroplasts to 3T3 cells(Figure 10J) and significantly promote their migration(Figure 10K). These results suggest that hybrids can effectively load chloroplasts and adapt to the wound microenvironment under light-driven conditions to facilitate healing.

### Hybrids loaded with C-dots can adapt to highly oxidative damage microenvironments

In addition to addressing epidermal injuries, cell therapy faces substantial challenges related to oxidative damage when functioning in the body’s deeper tissues^36^. Our research reveals that the viability of hybrids considerably diminishes as hydrogen peroxide concentrations increase, even though their resistance to hydrogen peroxide is markedly greater than that of BMSC, RAW264.7, and MPC5 cells(Figure 11A). Recently, quantum dots have attracted significant interest due to their remarkable capabilities in tracking cells and their biocompatibility^37^. Among these quantum dots, carbon quantum dot nanozymes are widely applied to improve cellular anti-inflammatory and antioxidant functions, thus mitigating conditions such as inflammatory bowel disease, acute lung injury, and sepsis^38^. We developed a novel C-dots nanozyme using methodologies previously described, which demonstrates both commendable superoxide dismutase enzymatic activity and fluorescence properties. The particle size measures about 3 nm, showing good dispersibility and a unique lattice structure (Figure 11B). It exhibits UV absorption peaks at 250 nm, 400-450 nm, and 650-700 nm, aligning with existing literature (Figure 11C). As the concentration of C-dots increased, the uptake rate of hybrids also continued to rise(Figure 11D-E). However, the hybrids could not transfer the C-dots they carried to the co-cultured MPC5 cells(Figure 11F). The results from the CCK-8 assay indicated that at concentrations of C-dots equal to or less than 100 μg/ml, there was no impact on the viability of hybrids(Figure 11G), meanwhile significantly lowering ROS levels and apoptosis rates in the presence of hydrogen peroxide (Figure H-I). These results imply that C-dot nanozymes may enhance the capacity of hybrids to adapt to tissue microenvironments prone to high levels of oxidative damage, thus safeguarding their therapeutic efficacy.

**Figure 11.**
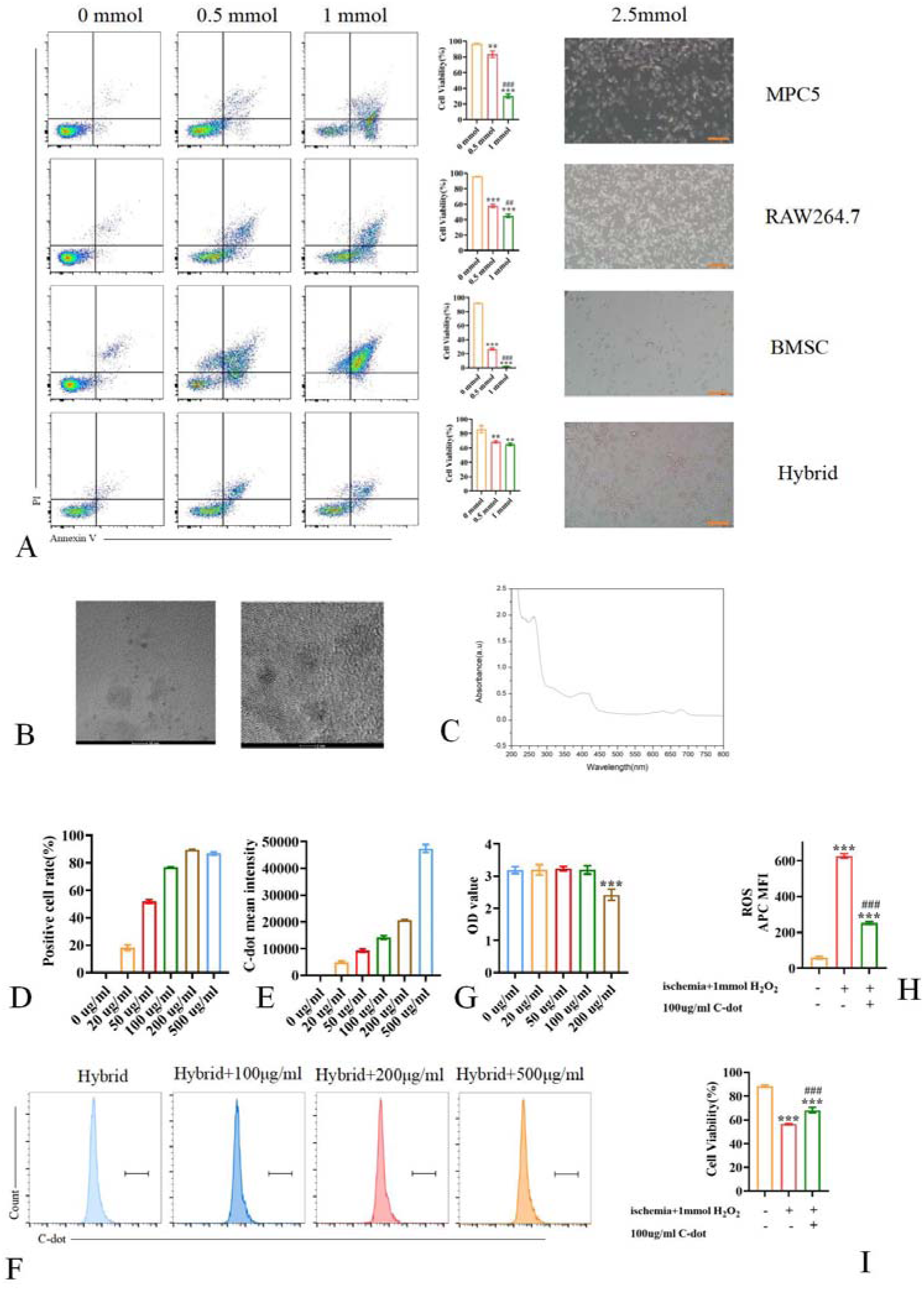
C-dots enhance the antioxidant damage resistance of hybrids. Flow cytometry and microscopic observation were used to detect the effects of different concentrations of H_2_O_2_ on the viability of MPC5, RAW264.7, BMSC, and hybrid(A). TEM analysis of C-dots(B). UV absorption peaks measurement of C-dots(C). Flow cytometry detection of the absorption rate of hybrids to C-dots(D-E), intercellular transmission of C-dots(F), and cell viability(G), ROS level(H), apoptosis(I) rate of hybrids. Data are represented as means±SD (n=3). ***P<0.001 vs 0 μg/ml in G(two-tailed t-test); ***P<0.001 vs no treatment group, ^###^P<0.001 vs only ischemia+1mmolH_2_O_2_ group in H and I(One-way ANOVA Tukey’s multiple comparison). Scale bar 200μm.

## Discussion

Both primary and secondary immune nephropathies result in the deposition of antibody immune complexes and the overactivation of immune cells, which ultimately leads to kidney damage. Most existing treatments primarily alleviate symptoms and slow disease progression, offering limited therapeutic benefits^39,40^. In recent years, cellular therapies such as CAR-T, M2/Mreg macrophage, and MSC therapy have demonstrated some efficacy by inhibiting antigen presentation and antibody production; however, they remain ineffective in clearing existing IgG immune complex deposits and promoting sustained kidney recovery^41–43^.

In this study, we propose the innovative BMSC-Macrophage hybrid therapy. Initially, during the construction of the hybrid, we utilized linker proteins Cov-19-S and hACE2, which were transiently transfected into BMSC and RAW264.7 cells, respectively, to induce cell fusion in vitro. In subsequent experiments, we observed that after culturing the hybrid for up to 72 hours, both proteins were no longer expressed, thus ensuring that they would not impact the results of the subsequent studies.

After successfully constructing the hybrid, we observed that the cytoplasm and cell membrane of BMSC and RAW264.7 fused, as demonstrated by fluorescence microscopy. Scanning electron microscopy revealed that the size and morphology of the hybrid were similar to those of RAW264.7, with no significant enlargement of the cells. However, the cell membrane exhibited increased wrinkling and bumpiness. Additionally, the glucose uptake rate of the hybrid was approximately 1.5 times higher than that of BMSC, as indicated by flow assay results. We hypothesized that some cytoplasm was shed during the fusion process, potentially alleviating the metabolic burden on the cells. Furthermore, the morphological changes in the hybrid membranes suggested that the cells could autonomously regulate the balance between the surface area of the cell membrane and the cell volume. This also indicated potential alterations in the distribution, interactions, and mechanical responses of receptors on the membranes. We did not further investigate mitochondrial division/fusion in hybrid or the metabolic state (aerobic glycolysis or oxidative phosphorylation) due to their sensitivity to external stimuli. In addition to regulating energy metabolism, mitochondria have been shown to influence MSC/macrophage differentiation and immunomodulation by interfering with autophagy, homeostasis, and communication between the nucleus, endoplasmic reticulum, lysosomes, and cytoskeleton^44^. Combined with the fact that the glucose uptake rate in the hybrid did not increase proportionally to the number of mitochondria, as described above, we hypothesize that the mitochondria in the hybrid are primarily responsible for a significant amount of intracellular communication tasks.

We employed superparamagnetic nanoparticles for purification, which are characterized by their low toxicity, ease of absorption, ability to be guided by magnetic fields, and suitability for nuclear magnetic imaging and in vivo tracking^25^. Previous studies have demonstrated that these nanoparticles enhance the antioxidant and pro-angiogenic capacities of MSCs and facilitate the differentiation of M1-type macrophages into M2-type macrophages^23^. Our observations indicated a reduction in ROS levels in nanoparticle-labeled hybrids prior to purification, suggesting that the superparamagnetic nanoparticles possess antioxidant properties in these hybrids. Following purification, we achieved a purification efficiency of approximately 90%. Furthermore, we found that the transfection reagent hindered the re-adhesion of RAW264.7 cells to the wells and effectively eliminated unfused macrophages. To optimize the purification process, we employed trypsin treatment, and flow cytometry analysis of single-stained cells revealed that a proportion of the cells lost fluorescence signals, leading to potential false negatives. Subsequently, we conducted long-term cultures of the purified cells and noted that as the culture duration increased, the presence of BMSCs became essentially undetectable, indicating that our purification method was both successful and efficient.

In our cloning experiments, we observed that BMSC did not readily form clonal clusters, which aligns with previous reports^45^. Although the cells utilized in our experiments were immortalized, it remains to be demonstrated whether the hybrid prepared from primary isolated BMSC and macrophages can undergo proliferation in subsequent experiments. Nonetheless, it has been reported that the fusion of primary glial cells and embryonic stem cells can result in the formation of clones^46,47^.

The qRT-PCR results indicated that hybrid CD16, CD32a, and CD64 expressions were significantly increased, suggesting that hybrids possess a strong capacity to bind IgG Fc segments. The expression of CD45 was higher than that of BMSCs but comparable to that of RAW264.7 cells, serving as a crucial reference point to regulate the excessive binding of hybrids to IgG. The binding of ITGAM to the FcγR receptor promotes phagocytosis and activates the ADCC effect. Although the expression of ITGAM in hybrids was upregulated, our investigation into the ADCC effect revealed that hybrids did not exhibit a significant ADCC effect. Furthermore, the levels of anti-inflammatory and antioxidant factors in hybrids were significantly elevated compared to both BMSCs and RAW264.7 cells, while the expression of adhesion factors ICAM-1 and CXCR4 was increased, facilitating the migration of syncytiotrophoblasts to the site of inflammatory injury. Additionally, our subsequent studies on IgG internalization and efferocytosis in hybrids demonstrated that anti-inflammatory and antioxidant factors were significantly increased. However, hybrids did not display a robust macrophage-like pro-inflammatory phenotype typical of M1 type cells and instead maintained high anti-inflammatory and antioxidant functions. In subsequent co-culture experiments involving hybrid cells with DOX-treated MPC5 cells, we observed that hybrid cells significantly promote the repair of MPC5 damage. Therefore, we suggest that hybrid cells are more inclined to exhibit anti-inflammatory properties and enhance damage repair when stimulated by IgG, ACs, and certain inflammatory factors. We will continue to investigate this further to validate our findings.

CD46, CD55, and CD59a are complement inhibitory factors that effectively inhibit complement-dependent cytotoxicity (CDC) ^48^. Compared to MPC5, CD46 and CD59a were significantly upregulated in the BMSC group, whereas only CD46 expression was upregulated in the hybrid group. This suggests that both BMSC and hybrid exhibit a certain inhibitory effect on complement activation; however, the hybrid group demonstrated a significant decrease in CD46, CD55, and CD59a levels compared to BMSC. It has been reported that maintaining lower levels of CD46, CD55, and CD59a during allogeneic infusion MSC therapy facilitates timely MSC clearance and reduces tumorigenicity^49,50^. Therefore, the hybrid may be safer for cell therapy compared to BMSC.

The IFNγ/IL-1β pre-activation regimen utilized in this study was directly referenced from existing research^51^. This regimen not only promotes the internalization of hybrid IgG and enhances the inhibitory effect of the hybrid on the proliferation of iDCs, but also increases the expression of PDL-1 to a certain degree. However, its long-term efficacy remains unknown, and we will subsequently explore more suitable hybrid pre-activation protocols.

We found that the hybrid can effectively clean IgG from the site of inflammatory injury. Although we observed that the hybrid has partial uptake of MPC5 cell membrane, the available experiments are insufficient to conclusively demonstrate that this process is efferocytosis. However, we favor this proposed pathway of action, which we plan to verify further. Similarly, while numerous reports indicate that MSCs promote renal injury repair through vesicle and mitochondrial delivery^52^, the specific mechanisms underlying the secretion of vesicle contents and mitochondrial delivery by the hybrid, as identified in the present study, remain unclear and warrant further investigation. In the study of efferocytosis, we employed the classical method for preparing ACs. Our findings indicate that, contrary to the previously reported macrophage phagocytosis of ACs^53^, hybrid exhibited a certain degree of saturation, resulting in a significant slowdown of proliferation. We speculate that this phenomenon may represent a self-protective mechanism that facilitates the adequate degradation of ACs while preventing an imbalance in own metabolism.

iDCs possess a robust antigen-presenting function, play a crucial role in both humoral and cellular immunity, and are also targets for anti-transplant rejection drugs^54^. Numerous studies have demonstrated that MSCs can inhibit the maturation, activation, and migration of iDCs, reduce the expression of inflammatory factors, and diminish the antigen-presenting capacity of iDCs^55^. In our study, we found that hybrid can effectively inhibit iDC proliferation and upregulate PDL-1 expression, which could hinder antigen presentation and reduces the production of new IgG. Furthermore, upon investigating the interaction mechanism between hybrid and iDCs, we were existed to discover that hybrid releases vesicles as it migrates. These vesicles are taken up by iDCs along the migratory path of hybrid, resembling the mechanism of migrasomes^56^. Although the morphology of these vesicles differs significantly from that of conventional migrasomes, subsequent pretreatment of hybrid with cell migration inhibitors revealed a significant attenuation in the uptake of hybrid vesicles by iDCs, suggesting that the vesicle delivery we observed may indeed belong to a type of migrasome. Recent studies have demonstrated that cardiomyocytes can expel dysfunctional mitochondria and other cellular components via large extracellular vesicles known as exophers (approximately 4μm in size)^57^. These exophers are taken up and cleared by cardiac-resident macrophages, thereby maintaining cardiomyocyte health and cardiac function. Furthermore, recent investigations have identified a vesicle-like structure called blebbisome, which can be autonomously released by both normal and tumor cells. These structures can reach a size of approximately 20 μm and exhibit notable expansion and contraction behaviors. Blebbisomes contain various cellular organelles and cytoskeletal components, yet lack nuclei. Remarkably, blebbisomes demonstrate a degree of ‘intelligence’, autonomously secreting and absorbing mitochondria and other components. They also mediate cellular communication and adapt to external environmental changes^58^. Although extracellular vesicles are currently classified into categories such as exosomes, microvesicles, apoptotic vesicles, exophers, and migrasomes based on size, surface markers, and contents, we propose that smaller vesicles may also possess functions similar to those of larger vesicles, and vice versa, influenced by environmental and stimulatory factors^59^. Additionally, we observed that vesicles delivered by hybrids to iDCs continue to be exchanged among DCs, thereby amplifying their influence. Moreover, it appears that the hybrid has the ability to autonomously recognize iDCs that have internalized its vesicles. This capability allows the hybrid to attract surrounding iDCs and recruit nearby hybrids to act in concert, which may be a manifestation of intelligence.

In the actual treatment process, utilizing primary cells can yield superior therapeutic effects; however, the high costs and prolonged culture periods associated with primary cells present significant challenges. Although the BMSC and RAW264.7 cells employed in our experiments are immortalized, they can be utilized to produce cell-derived nanovesicles, which offer a cost-effective, convenient, and readily accessible alternative. Increasingly, research has focused on the nanodecoy role of nanovesicles. Studies have reported that THP-1 and 293T cells can be engineered to create hybrid membranes with high expression levels of IL6R and TNFR on their surfaces. These membranes can serve as inflammatory factor nanodecoys, effectively mitigating inflammatory storms in infected pneumonia tissues^60^. Notably, the hybrid NV we produced exhibits high expression of CD16, CD64, PDL-1, and IL-6R, enabling it to effectively target and reside in IgG-deposited tissues while exerting immunosuppressive effects through PDL-1. Moreover, due to its elevated IL6R expression, it can effectively adsorb IL-6, making it a valuable adjunct to hybrid cell therapy. However, despite the high expression of IL-6R in our NV, the expression levels of the IL-1β receptor and TNF receptor remain low. Some researchers have employed chimeric receptor display technology to enhance the expression of various inflammatory factor receptors on the surfaces of NVs or exosome membranes^61^. We plan to incorporate this technology to optimize the hybrid NV and improve its efficiency in trapping inflammatory factors.

In this study, we innovatively explored the empowering effects of transplanting spinach chloroplasts on hybrids^62^. Previous reports suggested that chloroplasts needed to be encapsulated by biomembranes or similar materials to persist for extended periods in recipient cells, including macrophages^63^. However, the naked chloroplasts we extracted could effectively be retained in hybrids for over 24 hours, contradicting the literature which reported that chloroplasts would rupture and disperse throughout the mammalian receptor cells within 30 minutes^64^.

We speculate that the hybrid cells may inherit the non-degradative retention capabilities for xenobiotics/xenografts from MSCs, and that this retention effect could be facilitated through endogenous vesicle encapsulation^65^. A similar mechanism has recently been identified in Sacoglossans, specifically an organelle known as the Kleptosome, which is a membrane-enclosed symbiotic chloroplast capable of photosynthesis or direct decomposition to provide energy^66^.

In subsequent experiments, we observed that chloroplasts pre-labeled with green fluorescence were also present in 3T3 cells co-cultured without direct contact with the hybrids, suggesting that hybrids can transfer chloroplasts between cells via vesicular secretion. Furthermore, the fusion of MSCs with tumor cells or embryonic stem cells resulted in the retention of some organelles, chromosomes, or cytoplasmic components from non-MSCs^67^. This phenomenon was also observed in human-mouse hybrid cells, indicating that MSCs possess a unique compatibility with exogenous components. Additionally, it should be noted that the lighting conditions used in this study was referenced from some reports on plant cultivation^68^. For actual application in clinical treatment, further exploration is needed to develop protocols that can enhance patient compliance.

Another exploratory aspect of this study was to investigate whether hybrid could be loaded with c-dots to enhance its adaptability to high oxidative damage environments. Our findings indicate that, although previously reported c-dots can be absorbed by hybrid and significantly improve its antioxidant capacity, the required incubation dose for absorption is excessively high^69^. Moreover, the absorbed dots are not delivered to neighboring cells as vesicles, which contradicts existing work. Therefore, in subsequent research, we will also explore c-dots that are more suitable for absorption by our hybrid.

Of cause, hybrids can also function as viable drug carriers for the intelligent and controlled release of immunomodulatory drugs. This objective can be realised through a range of methodologies, including click chemistry, hydrogel micropatches and gene editing. Collectively, these approaches are poised to accelerate the development of novel therapeutic interventions for immune nephropathies.

## Supporting information

https://1drv.ms/w/c/58c6f3f948f90f44/ETcgOOxKj3JBi1ogxnC9hFYBQEqKbaYD7FYvFF-Go94Jxg?e=LM8W3T

https://1drv.ms/w/c/58c6f3f948f90f44/ETt9iibbWV1AqCpC4V9EeSMBcGO7K-Wixsch2NB5n7ye8w?e=0lDytF

## Acknowledge

This work was supported by awards from the Natural Science Foundation of Jiangsu Province (BK20231189), and PanFeng Innovative Team Project of the The Third Affiliated Hospital of Soochow University.

## Author Contributions

Jian Shi and Min Fan designed the experiments and data analysis. Jian Shi, Xiaotong Wu and Anqi Yang performed experiments, data analysis and wrote the manuscript. Jiawei Pan, Yangyang Sun, Jundong Zhu, Yuan Zhang and Linglong Jiang revised the manuscript. Min Fan supervised the project.

## Competing financial interests

The authors declare no competing financial interests.

## Notes

### Competing Interest Statement

The authors have declared no competing interest.

### Summary of Updates

After our inspection, we found that the term used was incorrect. We changed "BMSC-Macrophagy" in the title, abstract(objective), and main text to "BMSC-Macrophage". We will display the modified parts in the document in red letters. The spelling errors in the authors' names have been updated.

