## Supplementary material for "Construction, Phenotypic Characterization, and Immunomodulatory Function Study of BMSC-Macrophage Hybrid in vitro": https://1drv.ms/w/c/58c6f3f948f90f44/ETt9iibbWV1AqCpC4V9EeSMBcGO7K-Wixsch2NB5n7ye8w?e=0lDytF

**Table 1. The primer sequences used for qRT-PCR.**

| Primer | Sequence (5’-3′’) |
| --- | --- |
| CD32a | Forward:5’-TGCTAAATCTTGCTGCTGGGACTC-3’  Reverse:5’-GTCAGTGTCACCGTGTCTTCCTTG-3’ |
| CD16 | Forward:5’-CCAAGCCTGTCACCATCACTGTC-3’  Reverse:5’-AGGAGGCACATCACTAGGGAGAAAG-3’ |
| CD64 | Forward:5’-GGCTGCAGAGAGAGAAGAAATA-3’  Reverse:5’-CATCGCTTCTAACTTGCTGAAA-3’ |
| CD45 | Forward:5’-TGGATGGAGGCAAGCAGGATG-3’  Reverse:5’-ATTGGCAGCATGTTCTGGTTCC-3’ |
| ITGAM | Forward:5’-GCATCAATAGCCAGCCTCAGTG-3’  Reverse:5’-AGCCAGGTCCATCAAGCCATC-3’ |
| TSG-6 | Forward:5’-CTCACCTACGCCGAAGCCAAG-3’  Reverse:5’-CCATCCATCCAGCAGCACAGAC-3’ |
| SOD2 | Forward:5’-TGTTACAACTCAGGTCGCTCTTCAG-3’  Reverse:5’-CTTGATAGCCTCCAGCAACTCTCC-3’ |
| IDO | Forward:5’-GTGTGAATGGTCTGGTCTCTGTGAG-3’  Reverse:5’-ACATTTGAGGGCTCTTCCGACTTG-3’ |
| PGE-2 | Forward:5’-TCAGCGTTATCCTCAACCTCATTCG-3’  Reverse:5’-GGTCCGTCTCCTCTGCCATCG-3’ |
| CD206 | Forward:5’-TGTACGCAGTGGTTGGCAGTG-3’  Reverse:5’-GCTCTGATGATGGACTTCCTGGTAG- 3’ |
| Arg1 | Forward:5’-CTCCAAGCCAAAGTCCTTAGAG-3’  Reverse:5’-GGAGCTGTCATTAGGGACATCA-3 |
| IL-10 | Forward:5’-CTTACTGACTGGCATGAGGATCA-3’  Reverse:5’-GCAGCTCATGGAGCATGTGG- 3’ |
| CD274 | Forward:5’-GCCTCAGCACAGCAACTTCAG-3’  Reverse:5’-TTGTAGTCCGCACCACCGTAG-3’ |
| TGF-β1 | Forward:5’-CCACCTGCAAGACCATCGAC-3’  Reverse:5’-CTGGCGAGCCTTAGTTTGGAC-3’ |
| ICAM-1 | Forward:5’-TGCCTCTGAAGCTCGGATATACC-3’  Reverse:5’-CGAACTCCTCAGTCACCTCTACC-3’ |
| CXCR4 | Forward:5’-GTCCACGCCACCAACAGTCAG-3’  Reverse:5’-ACCACCATCCACAGGCTATCGG-3’ |
| CD46 | Forward:5’-GCAGTAGCATGGTGATCTGTAGTG-3’  Reverse:5’-GGAGGCTTGGTAGGATGAGTAGG-3’ |
| CD55 | Forward:5’-GCTGTCTCTGTTGCTGCTGTC-3’  Reverse:5’-GTATGCCACTTTGCTTTGCTCAG-3’ |
| CD59a | Forward:5’-GCTTCTGGCTGTGTTCTGTTCC-3’  Reverse:5’-TTGATACACTTGCATTCCGGCTAC-3’ |
| VEGFA | Forward:5’-CTCCGTAGTAGCCGTGGTCTG-3’  Reverse:5’-CCTCTCCTCTTCCTTCTCTTCCTC-3’ |
| FGF2 | Forward:5’-CCCACACGTCAAACTACAACTCC-3’  Reverse:5’-AGCAGCCGTCCATCTTCCTTC-3’ |
| PDGFB | Forward:5’-ACCAGCAGTTTGAGCAGTATTTCC-3’  Reverse:5’-GAGATGAAGTACAGGCAGCAGTG-3’ |
| βcatenin | Forward:5’-GCCATCTGTGCTCTTCGTCATC-3’  Reverse:5’-TAACCACAACAGGCAGTCCATAATG-3’ |
| Ang1 | Forward:5’-GGGAAGATGGAAGCCTGGATTTC-3’  Reverse:5’-GTACTGCCTCTGACTGGTTATTGC-3’ |
| HGF | Forward:5’-CAGTCAGCACCATCAAGGCAAGG-3’  Reverse:5’-ACCAGGAACAATGACACCAAGAACC-3’ |
| iNOS | Forward:5’-GAAGACAACTGGACAGGAACCTCAC-3’  Reverse:5’-AAATCCCGACTCTGGCATTCACAC-3’ |
| TNF-α | Forward:5’-CTGAACTTCGGGGTGATCGG-3’  Reverse:5’-GGCTTGTCACTCGAATTTTGAGA-3’ |
| IL-6 | Forward:5’-CTGCAAGAGACTTCCATCCAG-3’  Reverse:5’-AGTGGTATAGACAGGTCTGTTGG-3’ |
| β-actin | Forward:5’-GTCGTACCACAGGCATTGTGATGG-3’  Reverse:5’-GCAATGCCTGGGTACATGGTGG-3’ |
| PsbA | Forward:5’-TTGCGGTCAATAAGGTAGGG-3’  Reverse:5’-GTGTGCTTGGGAGTCCTTG-3’ |
